# Complex chemical signals dictate Ah receptor activation through the gut-lung axis

**DOI:** 10.1101/2023.02.22.529529

**Authors:** Fangcong Dong, Iain A. Murray, Andrew Annalora, Denise Coslo, Dhimant Desai, Krishne Gowda, Jian Yang, Dingbowen Wang, Imhoi Koo, Fuhua Hao, Shantu G. Amin, Andrew D. Patterson, Craig Marcus, Gary H. Perdew

## Abstract

The aryl hydrocarbon receptor (AHR) mediates intestinal barrier homeostasis. Many AHR ligands are also CYP1A1/1B1 substrates, which can result in the rapid clearance within the intestinal tract, limiting AHR activation. This led us to the hypothesis that there are dietary substrates of CYP1A1/1B1 that increase the half-life of potent AHR ligands. We examined the potential of urolithin A (UroA) as a CYP1A1/1B1 substrate to enhance AHR activity in vivo. UroA is a competitive substrate for CYP1A1/1B1 in an in vitro competition assay. A broccoli-containing diet promotes the gastric formation of the potent hydrophobic AHR ligand and CYP1A1/1B1 substrate, 5,11-dihydroindolo[3,2-b]carbazole (ICZ). Dietary exposure to UroA in a broccoli diet led to a coordinated increase in duodenal, cardiac, and pulmonary AHR activity, but no increase in activity in liver. Thus, CYP1A1 dietary competitive substrates can lead to intestinal “escape”, likely through the lymphatic system, increasing AHR activation in key barrier tissues.

## INTRODUCTION

The aryl hydrocarbon receptor (AHR) is the only member of the basic-helix-loop/helix Per-Arnt-Sim family of proteins that is directly activated by low molecular weight molecules.^1^ Upon binding an agonist, the AHR undergoes a conformational change that results in translocation of the AHR bound to a heat shock protein complex into the nucleus.^2^ In the nucleus, the AHR complex encounters ARNT and forms a heterodimer capable of binding to dioxin-response elements in various promoter regions, resulting in altered transcription. The first identified AHR target gene was *CYP1A1* that encodes the cytochrome P450 1A1 protein and its expression is largely dependent on AHR activation. What followed was the examination of the AHR as a xenobiotic receptor that responds to the presence of halogenated polycyclic aromatic hydrocarbons, such as the tumor promoter 2,3,7,8-tetrachlorodibenzo-*p*-dioxin (TCDD). More recent studies, with the aid of *Ahr^-/-^* mice, have shown that the AHR is involved in a myriad of physiologic responses, including regulating immune and epithelial cell differentiation responses in barrier tissues, barrier protein production, and reproductive success.^3–7^ Importantly, AHR ligands can attenuate chemical and microbial insults, suggesting a possible therapeutic potential.^8–10^ One particularly promising study illustrated that dietary supplementation with a naturally occurring precursor of an AHR ligand, indole-3-carbinol, significantly diminishes disease caused by *Clostridium difficile* infection.^11^

Dietary plant phytochemicals are a source of AHR ligands within the intestinal tract. Flavones are perhaps the most abundant source of dietary AHR ligands. Flavones largely exist in plants conjugated with a carbohydrate moiety, that are cleaved by bacterial metabolism in the colon. A number of flavones occur naturally that vary in the number and position of hydroxy groups, including apigenin, quercetin, isorhamnetin, and galangin.^12^ All of these flavones have been shown to be either weak agonists or antagonists for the AHR. Perhaps the most potent source of dietary AHR agonists is through the consumption of cruciferous vegetables, which contain indologlucosinolates. Upon disruption of plant tissue, this phytochemical enzymatically breaks down into indole-3-carbinol, which undergoes acid condensation in the stomach. A variety of metabolites are formed, with the most potent AHR agonist being indolo[3,2-*b*]carbazole (ICZ), other metabolites are either not AHR ligands or are very weak agonist.^13^ The consumption of broccoli in a rodent diet at 15% (w/w) leads to activation of the AHR and subsequent induction of *Cyp1a1* in the intestinal epithelium. Furthermore, the presence of broccoli in the diet attenuates dextran sodium sulfate mediated colonic injury in an AHR-dependent manner.^14^ This underscores the potential for dietary sources of AHR ligands to modulate intestinal health. Urolithins have been identified as dietary antagonists for the human AHR and inhibitors or substrates for CYP1A1/1B1 enzymatic activity.^15, 16^ In contrast, urolithins appear to be very weak activators for the mouse AHR, and the mechanism of this activation has not been determined.^16^ Urolithins are generated through gut microbial metabolism of ellagitannins (ETs) and ellagic acid (EA). These phytochemicals are found in a variety of fruits and nuts, such as walnuts, raspberries, pomegranates and strawberries.^17, 18^ Because of the absorption characteristics of the various products of ETs, urolithins are the likely source of the various pharmacologic properties attributed to consumption of ETs.^18^

Several AHR ligands are reported to be metabolized by CYP1A1/1B1 to hydroxylated metabolites that can then undergo conjugation to generate a more polar metabolite that can be effectively eliminated. Perhaps the most studied example of this are polycyclic aromatic hydrocarbons, such as the carcinogen, benzo(a)pyrene.^19^ More recently, the endogenous AHR ligand 6-formylindolo[3,2b]carbazole (FICZ) also undergoes this same metabolic fate.^20^ This ability of AHR activation leading to the metabolism of AHR ligands results in a negative feedback loop that down-regulates AHR activity. Interestingly, cultured cells that essentially do not express the AHR can accumulate AHR ligands upon transient expression of the AHR. While co-expression of CYP1A1 and AHR leads to down-regulation of AHR activity.^21^ There is compelling evidence that this negative feedback loop occurs in the intestinal epithelium, which can lead to the prevention of intestinal AHR activity.^22^ Indeed, constitutive overexpression of CYP1A1 in the intestinal epithelium leads to ablation of AHR transcriptional activity, indicating that AHR ligands are present in the intestinal tract that are also CYP1A1 substrates. Thus, one possible mechanism of enhanced AHR activation is through the presence of competitive substrates or inhibitors of CYP1A1/1B1 that attenuates the metabolism of a given AHR ligand within the intestinal tract, leading to systemic AHR activation. However, whether this occurs in vivo due to the presence of dietary CYP1A1/1B1 inhibitors/substrates has not been established. In this study we determined that urolithins can act as competitive CYP1A1 substrates, increasing the half-life of the potent AHR ligand 5,11-dihydroindolo-[3,2-b]-carbazole (ICZ), which enhances AHR activation in the small intestine, thus allowing “escape” of an AHR ligand from the intestinal tract and subsequent activation of the AHR in the lung, establishing a gut-lung axis.

## RESULTS

### Dietary phytochemicals are required for both intestinal and pulmonary AHR activity

*Ahr^+/+^* mice were fed either a standard chow or semi-purified AIN93G diet and the level of duodenum, lung, and liver *Cyp1a1* expression was determined (Fig. 1A). *Cyp1a1* expression was just above the level of detection in mice on a semi-purified diet, indicating a lack of intestinal and lung AHR activity in the context of a diet devoid of phytochemicals. In contrast, a significant level of duodenal and pulmonary *Cyp1a1* expression was observed in mice on a chow diet. *Cyp1a1* was not detected in *Ahr^-/-^* duodenal and pulmonary tissues, indicating the requirement for AHR expression to mediate *Cyp1a1* expression, thus confirming that *Cyp1a1* is an appropriate marker of AHR activity.

**Figure 1.**
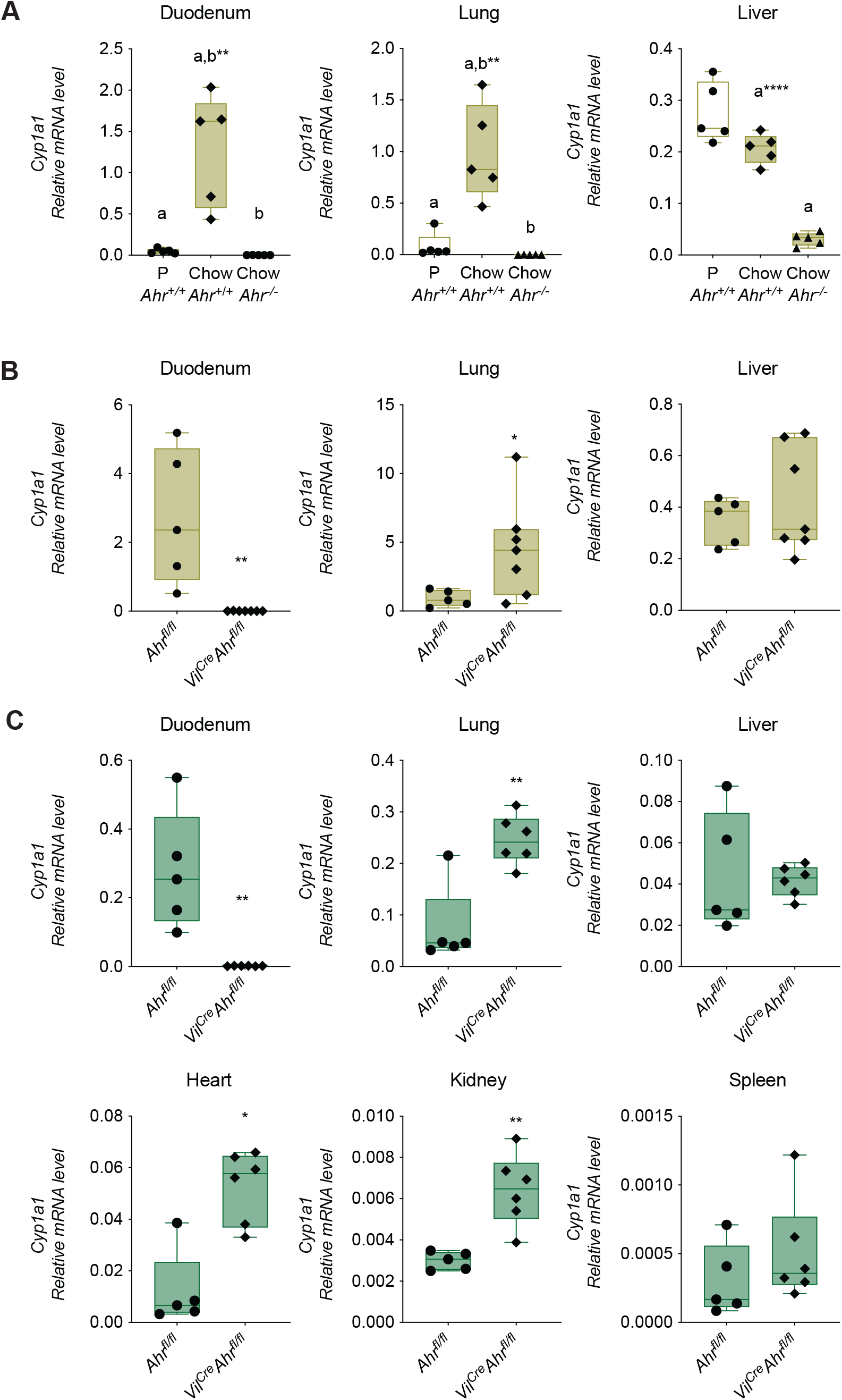
Chow diet leads to pulmonary *Cyp1a1* gene expression enhanced by the absence of intestinal epithelial AHR expression. (A) Duodenal, pulmonary, and hepatic *Cyp1a1* expression in *Ahr^+/+^* mice fed either a chow or semi-purified diet were compared to *Ahr^-/-^* mice fed a chow diet. (B) Duodenal, pulmonary, and hepatic *Cyp1a1* expression in *Ahr^fl/fl^* and *Villin^Cre^Ahr^bfl/fl^* mice on a chow diet. (C) Tissue-specific expression of *Cyp1a1* in *Ahr^fl/fl^* and *Villin^Cre^Ahr^fl/fl^* mice on a 15% broccoli diet. Statistical difference was analyzed using one-way ANOVA with Tukey’s multiple comparisons (A) and two-tailed unpaired Student’s t tests (B and C). Data are presented as mean + SEM (error bars). *p< 0.05, **p< 0.01, ****p<0.0001.

### Lack of AHR expression in the intestinal epithelium increases pulmonary Cyp1a1 expression

The level of *Cyp1a1* expression was determined in control *Ahr^fl/fl^* and enterocyte specific conditional *Ahr* knockout *Villin^Cre^Ahr^fl/fl^* mice fed a chow diet. *Cyp1a1* expression was not detected in the duodenum of mice that lack enterocyte AHR expression, indicating that AHR-mediated *Cyp1a1* expression is largely restricted to duodenal enterocytes (Fig. 1B). In lung an increase in *Cyp1a1* was observed in lung of *Villin^Cre^Ahr^fl/fl^* compared to in *Ahr^fl/fl^* mice, while no significant difference in *Cyp1a1* expression was observed in liver. This would support the concept that an AHR agonist that “escapes” CYP1A1 metabolism in the intestinal tract can lead to tissue-specific extra-intestinal AHR activation. One hypothesis that could explain these results is that an AHR agonist enters the lymphatic system, however the identity of AHR agonist in chow is not known. Therefore, a broccoli diet was utilized, which leads to the gastric formation of ICZ promoting AHR activation.^14^ ICZ is a potent hydrophobic planar ligand that would likely associate with lipid micelles within an aqueous environment. A 10% broccoli diet was fed to *Ahr^fl/fl^* and *Villin^Cre^Ahr^fl/fl^* for 3 days and tissues were isolated. The level of *Cyp1a1* expression was determined and the lack of AHR expression in the intestinal tract led to enhanced *Cyp1a1* expression in the lung (5-fold), heart (7-fold), and kidney (2-fold) (Fig. 1C). The observed increase in cardiac *Cyp1a1* observed in *Villin^Cre^Ahr^fl/fl^* is consistent with lymphatic drainage of the thoracic duct, which then enters the heart (Fig. 1C). In contrast, there was no significant increase in of *Cyp1a1* in liver and spleen.

### Urolithins are CYP1A1 inhibitors or competitive substrates

Several reports in the literature suggest that the presence of CYP1A1/1B1 substrates/inhibitors could increase the half-life of AHR ligands that are metabolized by CYP1A1/1B1. However, whether common dietary phytochemicals can modulate AHR activity in vivo through modulating CYP1A1/1B1 metabolism has not been firmly established. In our previous studies, urolithin A (UroA) in mouse hepatoma cells appeared to be a very weak activator of the mouse AHR, although the mechanism of activation was not determined.^16^ In addition, as tricyclic compounds, urolithins could be CYP1A1 substrates. Using an in vitro microsomal CYP1A1/1B1 assay system, the ability of six urolithins to inhibit metabolism of luciferin-CEE, a specific CYP1A1 substrate, was examined (Fig.2A). To have a significant level of CYP1A1 enzyme, cells were pretreated for 24 h with the prototypical AHR agonist TCDD prior to isolating microsomes. Critically, despite its status as an AHR ligand, TCDD is neither an inhibitor nor substrate for CYP1A1. The expression of microsomal CYP1A1 was confirmed by WES protein analysis in both Caco2 and Hepa 1 microsome fractions (Fig. S1). At 5 µM all urolithins tested were capable of attenuating CYP1A1 mediated metabolism of luciferin-CEE, with UroA, B, C and D exhibiting the most potent activity. Urolithin A, perhaps the most abundant urolithin, could inhibit CYP1A1/1B1 metabolism in both the Caco2 (human) and Hepa 1 (mouse) microsomal assay system, yielding similar IC_50_ values (Fig. 2B, C). To provide context to the relative potency of UroA as a competitive substrate/inhibitor of CYP1A1/1B1, UroA was compared to the known CYP1A1 substrate α-naphthoflavone (α-NF), and UroA exhibited ∼9-fold greater potency (Fig.S2). LC-MS/MS analysis confirmed that microsomal assays metabolize luciferin-CEE to luciferin (Fig. S3).

**Figure 2.**
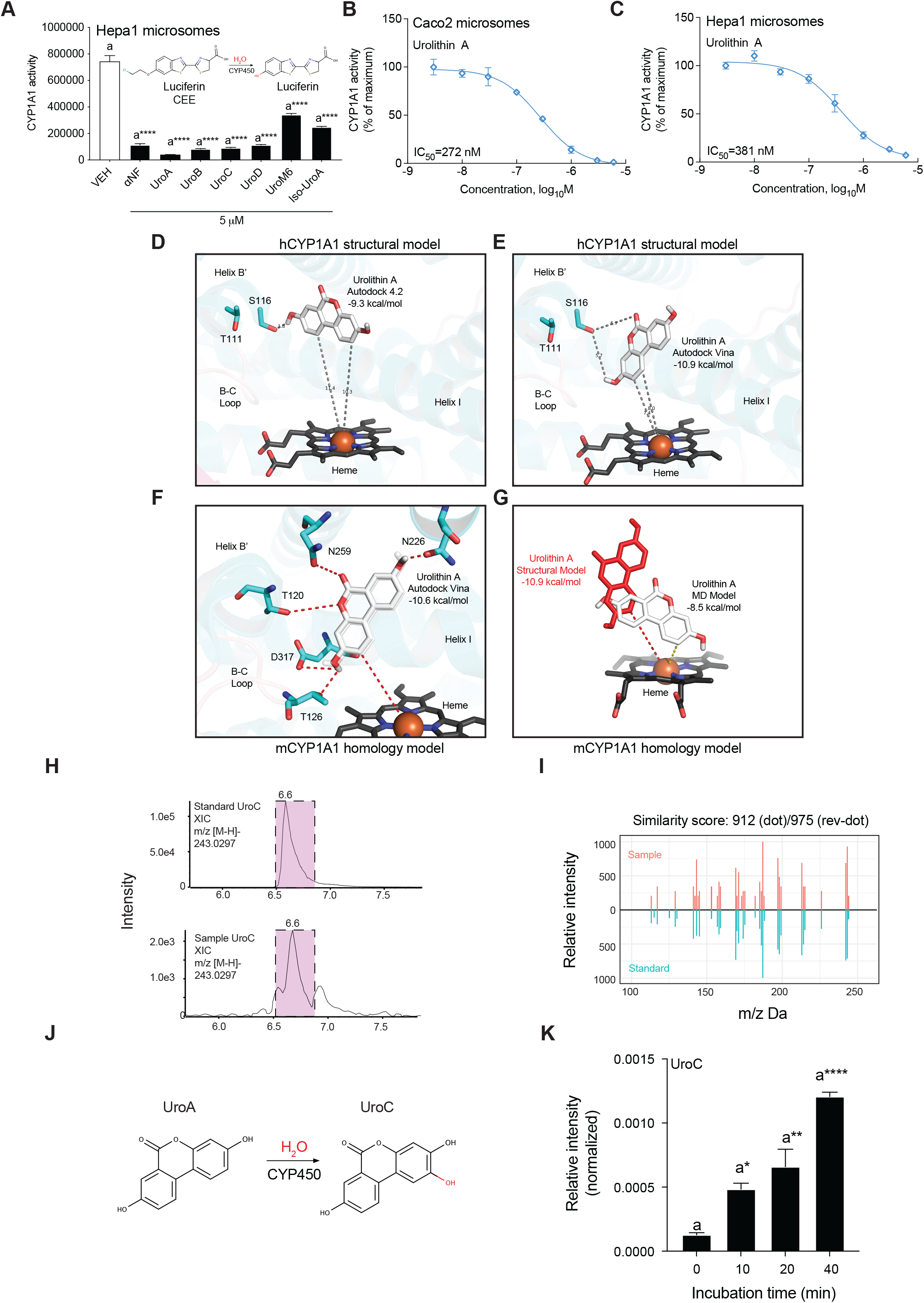
Urolithins are CYP1A1 substrates/inhibitors. (A) Six urolithins were co-incubated with luciferin-CEE CYP1A1 substrate in a Hepa 1 microsomal assay and compared to α-NF. (B, C) Dose-dependent inhibition of luciferin-CEE metabolism by UroA in the Hepa 1 or Caco2 microsomal assay system. (D) UroA binding properties in CYP1A1 were studied using both Autodock 4.2 and Autodock Vina. Low-energy Autodock 4.2 poses show UroA binds human CYP1A1 (hCYP1A1) with nanomolar affinity (hCYP1A1: 151 nM, -9.3 kcal/mol) in an allosteric site near the top of the active site positioning the C2 carbon 10.3 Å from the heme center. Similar results for UroA binding in a mouse CYP1A1 (mCYP1A1) homology model were obtained (mCYP1A1: 191 nM, -9.17 kcal/mol). Interactions between UroA and species-specific, active site residues Threonine-111 (T111) and Serine-116 (S116) are shown. (E) When UroA binding was studied using a more advanced program, Autodock Vina, a lower energy docking pose was identified (hCYP1A1: 17 nM; -10.9 kcal/mol) that positions the C2 carbon 7.4 Å from the heme center. Autodock Vina identified an alternate interaction between UroA and the B’ helix residue S116 in hCYP1A1, as shown. (F) Comparative Autodock Vina analysis in the mCYP1A1 homology model identified the same, low energy pose for UroA observed in the hCYP1A1 model shown in Figure 1B, positioning the substrate ∼3.0 Å closer to the heme center. Here, the C2 target carbon remains closest to the heme iron (7.6 Å) but the absolute binding energy was reduced (mCYP1A1: 191 nM, -10.6 kcal/mol. UroA (from mCYP1A1; white) is shown superimposed on the related result obtained from the hCYP1A1 structural model (in transparent white); minor repositioning of hydroxyl groups can be observed between the two nearly identical solutions. (G) An unbiased, MD simulation-based model of mouse CYP1A1 was also prepared to remove crystallographic artifacts of α-naphthaflavone binding from our homology model. After relaxation of the backbone and energy minimization of the substrate-free active site, UroA was able to dock within 4.6 Å from the heme center, well-positioned for target-specific oxidation of the C2 carbon. Reduced binding affinity was predicted for UroA in the unbiased MD model (-8.5 kcal/mol; mCYP1A1; Autodock Vina), but similar structural features of active site recognition were highlighted using both structure- and MD-simulated models of the CYP1A1 active site. Microsomes isolated from TCDD-treated Hepa1 cells were utilized an in vitro CYP1A1/1B1 metabolism assay. (H) Representative selected ion chromatogram (XIC) of UroC compared to authentic standards; (I) MS/MS spectral similarity plot of authentic UroC compared to putative UroC generated in microsomal assays; (J) Structural depiction of reaction route of UroC produced from UroA; (K) UroC was quantitated by LC-MS/MS from UroA/microsomal incubations in a time-dependent manner. Statistical difference was analyzed using one-way ANOVA with Tukey’s multiple comparisons (D). Data are presented as mean + SEM (error bars). *p< 0.05, **p< 0.01, ****p<0.0001.

### Predictive molecular modeling of UroA metabolism by CYP1A1 orthologs

Next, we wanted to test whether UroA is an inhibitor or a substrate for CYP1A1. To guide the search for a possible CYP1A1-dependent metabolite of UroA, we utilized a computational docking approach. The substrate binding properties of UroA were studied within the active site cavity of both human and mouse cytochrome P450 1A1 (CYP1A1) using Autodock 4.2 and Autodock Vina. A homology model for mouse CYP1A1 (mCYP1A1) was developed using the crystal structure of human CYP1A1 (PDB: 4I8V; see methods). Structure-based models of both human and mouse orthologs were used to compare species-specific differences in UroA substrate recognition. MD-simulation was also utilized to reduce crystallographic bias in the CYP1A1 active site, as needed to dock UroA in close proximity to the heme center (see methods). Primary amino acid sequence alignments, substrate recognition sequence (SRS) motif and structural analysis of mouse and human CYP1A1 have also been provided for additional context in Figure S4. Using Autodock 4.2, we calculated that UroA binds mouse and human CYP1A1 with nanomolar affinity (hCYP1A1: 152 nM; -9.3 kcal/mol and mCYP1A1: 191 nM; -9.17 kcal/mol; see Table S1), within an allosteric binding site, above the heme center (Fig. 2D). An alternative, lower energy binding solution for UroA, in the hCYP1A1 model, was identified using the related program Autodock Vina (Fig. 2E), which docked the substrate ∼2.0 Å closer to the heme center than Autodock 4.2. Comparative docking analysis performed with the mCYP1A1 homology model revealed a similar docking pose for UroA, harmonizing docking results for both mouse and human models of CYP1A1 (Fig. 3F). Autodock Vina also predicted nearly identical affinity with respect to the dissociation binding constants (K_D_) for Urolithin A in both mouse and human models of CYP1A1 (hCYP1A1: 17 nM; -10.9 kcal/mol; mCYP1A1: 10 nM; -10.6 kcal/mol; see Table S1). No Autodock 4.2 or Vina-based docking solutions positioned the putative C2 target carbon of Urolithin A closer than 7.4 Å to the heme iron, the likely site for hydroxylation. To explore whether substrate and crystal-lattice conformational bias in the crystal structure of CYP1A1 (*PDB: 4I8V, copy A*) constrained UroA docking to the observed allosteric pocket, we performed an MD-simulation based energy minimization and simulated annealing of our mouse CYP1A1 homology model, in the absence of the crystallographic substrate α-NF. As shown in Figure 3G, stripped of native electrostatic interaction biased for α-NF binding, UroA was free to bind deeper into the mouse CYP1A1 active site and position the target C2 carbon in a catalytically favorable position within 4.6 Å of the heme center. While this pose is most predictive of known substrate metabolism, there was significant trade-off in UroA binding energy deeper in the MD-simulated pocket (578 nM; -8.5 kcal/mol; Autodock Vina) as substrate affinity was reduced ∼30-50 fold compared to the crystal structure-based models of mouse and human CYP1A1. These results suggest that the most likely hydroxylated UroA product would be UroC.

**Figure 3.**
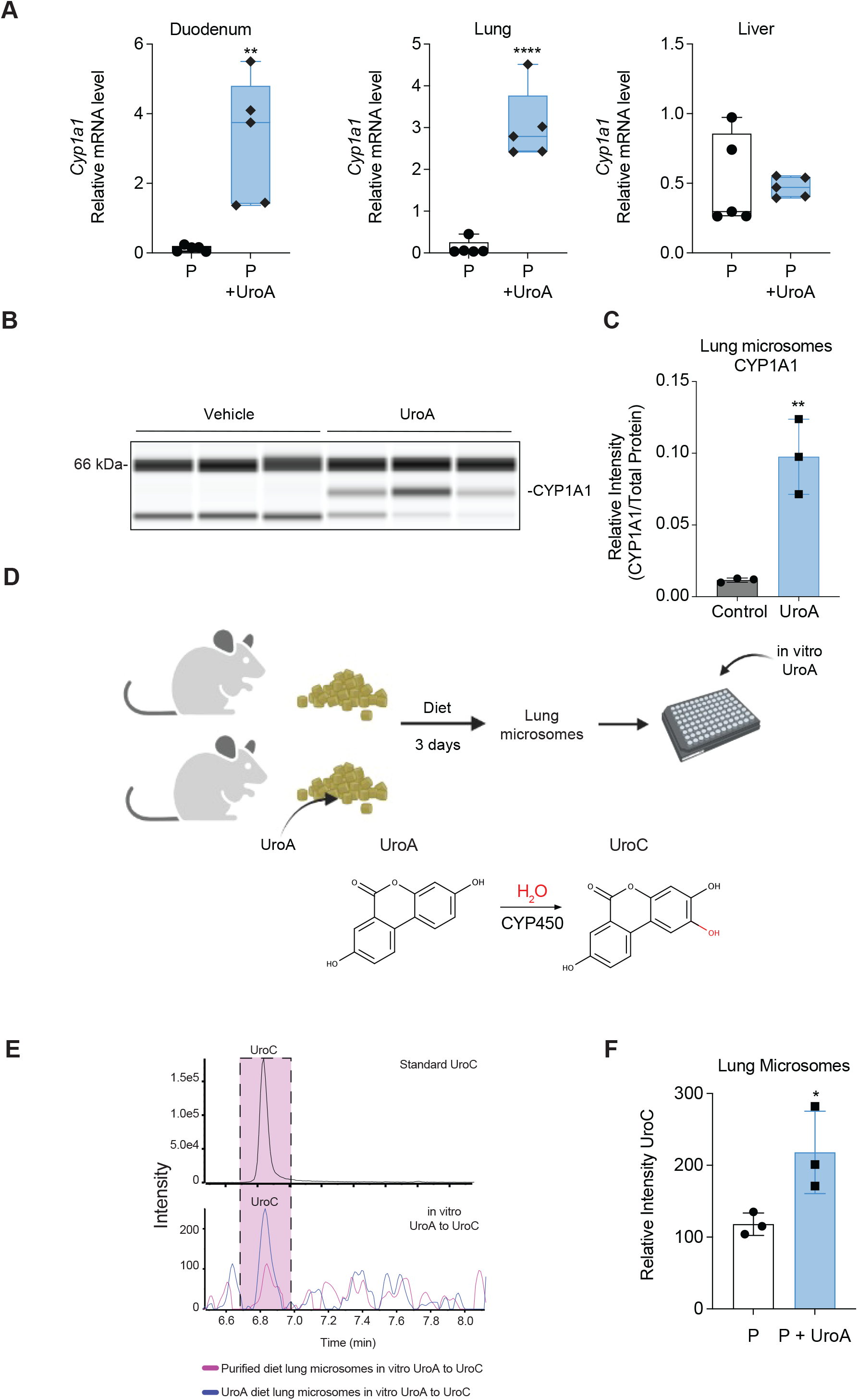
Dietary UroA increases pulmonary CYP1A1 expression. (A) Mice were fed a semi-purified diet for one week followed by 3 days on either a semi-purified diet or the same diet + UroA. The level of C*yp1a1* expression was determined in duodenal, pulmonary, and hepatic tissues. (B) Lung microsomes were isolated from lungs fed the same diets in panel A and the level of CYP1A1 examined using the WES system. (C) Quantitation of the WES data in panel B. (D) Scheme for in vitro microsomal assay. (E, F) Representative LC-MS/MS analysis of UroC levels after UroA incubation with lung microsomes described in panel B. Statistical difference was analyzed using two-tailed unpaired Student’s t tests (A,B and F). Data are presented as mean + SEM (error bars). *p< 0.05, **p< 0.01, ****p<0.0001.

### UroA undergoes microsomal metabolism to UroC

Hepa 1 microsomal metabolism assays containing UroA were extracted and analyzed by LC-MS/MS. After incubation with microsomes a m/z 243.0297 peak was observed at 6.6 min, identical to the UroC standard (Fig. 2H). Comparison of the MS/MS spectra for the 6.6 min peak from the microsomal incubation and authentic standard of UroC revealed a similar fragmentation pattern, which confirms that UroA is hydroxylated by CYP1A1 to form UroC with a dot and reverse spectrum similarity scores calculated as 912.5 (dot) and 975.4 (reverse-dot) out of 1,000, respectively (Fig. 2I and S5).^23^ In addition, incubation of UroA with Hepa 1 microsomes resulted in a time-dependent increase in UroC (Fig. 2J, K). This data establishes that UroA is a CYP1A1 substrate and may effectively compete with other CYP1A1 substrates.

### Dietary UroA promotes intestinal and extra-intestinal CYP1A1 expression and microsomal activity

To explore the influence of dietary UroA on AHR activation potential, mice were fed UroA for 3 days in a semi-purified diet (Fig. 3A). UroA consumption led to a similar marked increase in *Cyp1a1* expression in the duodenum and lung, but no significant difference was observed in liver. Lung microsomes were isolated from mice fed either a semi-purified or semi-purified with UroA diet, subjected to WES protein analysis of CYP1A1 levels and normalized to total protein (Fig. 3B, Fig. S6A). The addition of UroA to the diet resulted in a 9-fold increase in CYP1A1 protein levels (Fig. 3C). Next, the ability of lung microsomes to metabolize UroA to UroC was examined in an in vitro microsomal P450 metabolism assay. Lung microsomes from mice fed UroA demonstrated an increased capacity to metabolize UroA to UroC (Fig. 3D-F).

### Bacteria, dietary indole-3-carbinol and broccoli fail to influence the tissue-specific pattern of AHR activation

To test whether bacteria either in the lung or gut play a role in these observations, germ-free mice on a semi-purified diet were switched to the same diet with UroA for 3 days and results were similar to those observed in conventional mice in figure 3A (Fig. 4A). This would suggest that microorganisms in the gut do not affect the pattern of activation. Interestingly mice fed a semi-purified diet with indole-3-carbinol, which undergoes acid condensation to ICZ in the stomach, revealed *Cyp1a1* induction in the duodenum and lung, but not in the liver. To provide relevance to these observations and allow the examination of a possible mechanism of UroA mediated increase in AHR activation, mice were fed a semi-purified diet with 10% broccoli for one week followed by 3 days with or without UroA. The level of *Cyp1a1* expression was examined in anatomical segments of the intestinal tract, and a significant increase in *Cyp1a1* expression was observed in the duodenum, jejunum, ileum and lung when fed UroA in the broccoli diet (Fig. 4B, C). In contrast, the presence of UroA in a broccoli diet failed to alter *Cyp1a1* expression in the colon, spleen, or liver. The activation of *Cyp1a1* from broccoli consumption is due to the in-situ production of ICZ in the stomach. The concentration of UroA in portal blood from broccoli and UroA fed mice was assessed by LC-MS/MS analysis and determined to have mean concentration of ∼300 nM (Fig. S6B), a level that is unlikely to directly influence AHR activation as a ligand.

**Figure 4.**
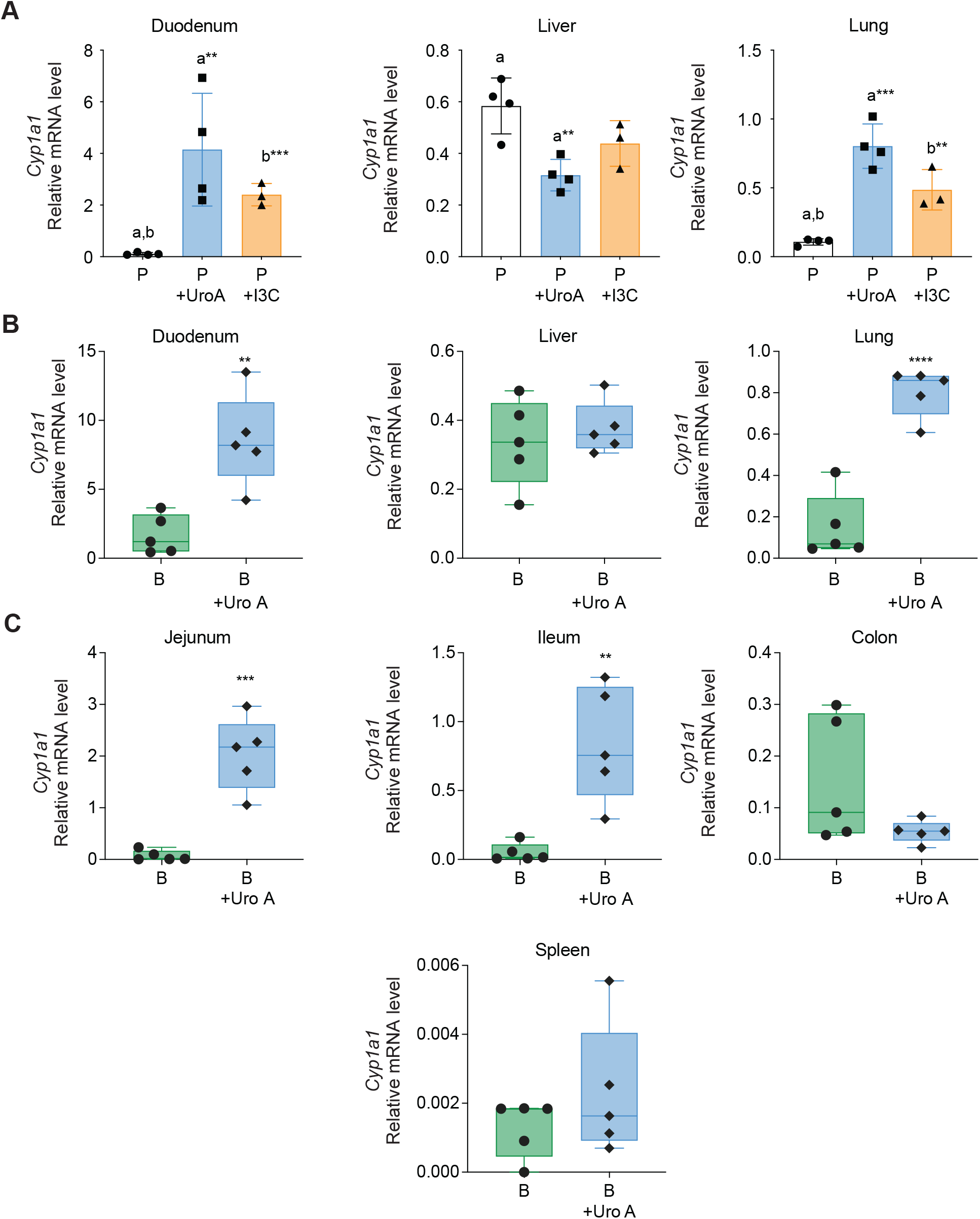
Tissue specific modulation of *Cyp1a1* by UroA and broccoli. (A) Germ-free mice were fed a semi-purified diet with or without 4 mg of UroA/g of food or 125 µg of indole-3-carbinol/g of food for 3 days and *Cyp1a1* levels determined. (B, C) Mice were fed a semi-purified diet for one week followed by semi-purified diet + 10% broccoli or semi-purified diet + 10% broccoli + 4 mg UroA/g of food for 3 days and *Cyp1a1* measured in the indicated tissues. Statistical difference was analyzed using one-way ANOVA with Tukey’s multiple comparisons (A) and two-tailed unpaired Student’s t tests (B). Data are presented as mean + SEM (error bars). **p< 0.01, ***p<0.001, ****p<0.0001.

### ICZ inhibits CYP1A1/1B1 metabolism in vitro

ICZ was added to the CYP1A1/1B1 mouse microsomal assay system and inhibited luciferin-CEE substrate metabolism at an IC_50_ of 616 nM (Fig. S7A-C). Next. we wanted to determine whether ICZ is a CYP1A1/1B1 substrate or inhibitor.

### Predictive molecular modeling of CYP1A1 metabolism of ICZ

ICZ is a hydrophobic molecule produced in the stomach upon consumption of broccoli that has limited solubility in an aqueous microsomal assay system. Therefore, to gain a better understanding of relative differences in the CYP1A1/1B1 metabolic turnover of ICZ and UroA, we turned to molecular modeling utilizing the crystal structures of the CYP1A1/1B1 catalytic domain. The nature of competitive binding between UroA, and the AHR ligand ICZ, was explored using the docking stimulations. ICZ binds hCYP1A1 (-11 kcal/mol; 8 nM) and mCYP1A1 (-11.3 kcal/mol; 84 nM) models in a common pose with higher affinity than UroA in any conformation, when using Autodock 4.2 (Fig. 5A; Table S1). Similar docking poses for ICZ were obtained using Autodock Vina, but picomolar level binding affinity was predicted (hCYP1A1: 49 pM; -14.1 kcal/mol; mouse (135 pM; -13.5 kcal/mol). In CYP1A1, hydrophobic interactions between ICZ and conserved β1-4 loop residue Valine-386 (V386) and β4-1/4-2 loop residue Theonine-501 (T501) regulate the docking regiospecificity of ICZ. ICZ binds CYP1A1 in a single robust conformation that positions the C2 carbon 4.5 Å from the heme center. Here, ICZ would superimpose on the docking coordinates of the two, lowest energy UroA docking solutions obtained with structure-based models of CYP1A1 (Fig. 5B). This observation, paired with the picomolar affinity, strongly imply that ICZ will consistently outcompete UroA for binding in CYP1A1’s proximal active site, potentially inhibiting its co-metabolism.

**Figure 5.**
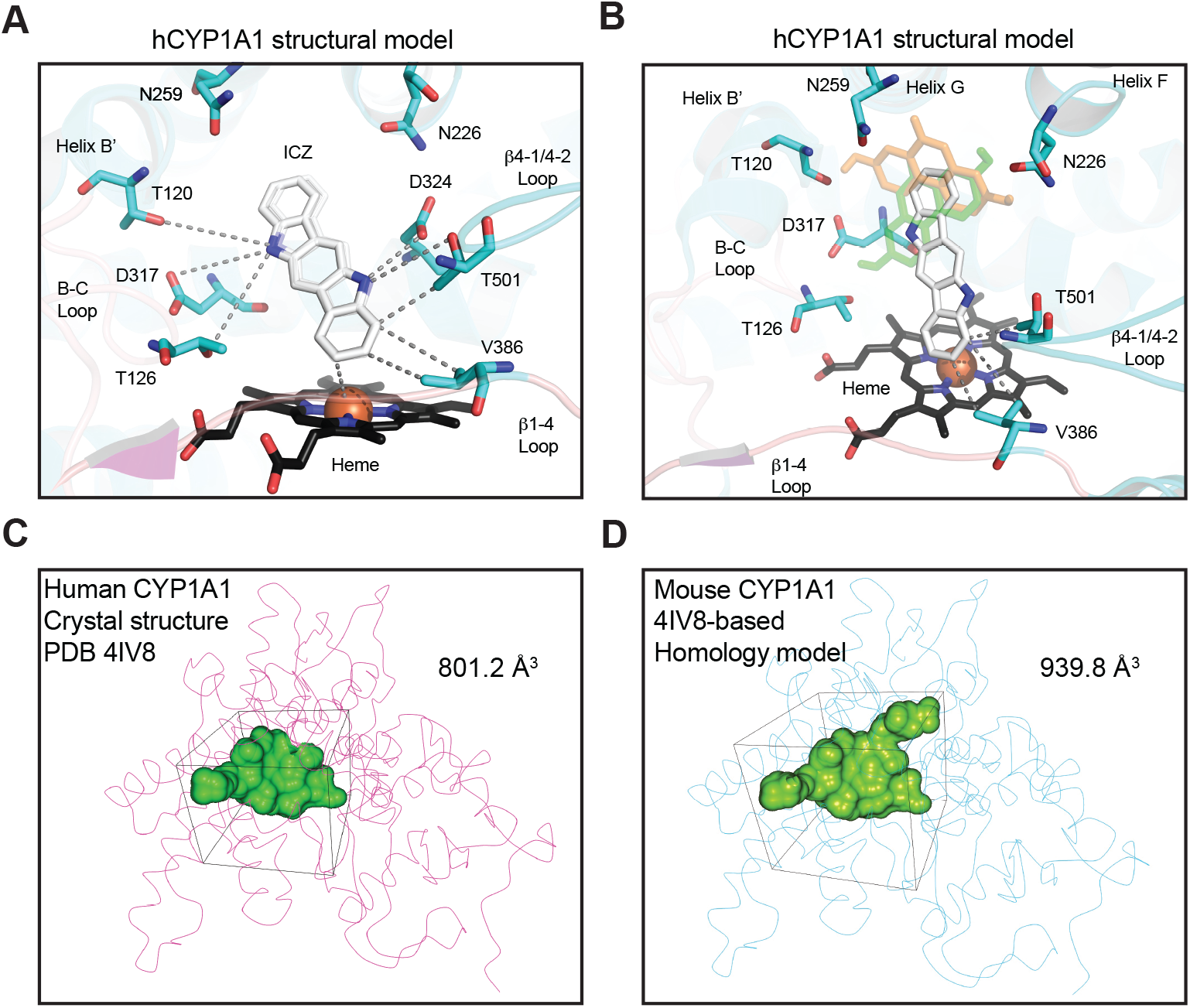
ICZ and UroA competitive docking models of human CYP1A1. (A) The nature of ICZ binding to the human CYP1A1 substrate pocket was explored using Autodock 4.2 and Vina with a very consistent conformation that positions the C2 carbon 4.5 Å from the heme center. (B) ICZ superimposed over both low and high energy docking positions for Urolithin A (shown in transparent orange and green sticks). (C) Our computational docking model for human CYP1A1 was derived from crystal structural coordinates (PDB: 4I8V; copy A). Using CAVER 3.0, we calculated a square active site cavity for human CYP1A1 of 801.2 Å^3^. (D) A homology model for mouse CYP1A1 was developed using the human crystal structure and the SWISS-MODEL server. Mouse CYP1A1 has a predicted active site volume 138.6 Å^3^ larger than the human isoform due primarily to species-specific differences in the SRS-3 region (see Figure S3D).

Overall, ICZ is predicted to bind the CYP1A1 active site with sub-nanomolar affinity, in a robust configuration that would allow it to compete for substrate access with virtually any CYP1A1 substrate, including more soluble compounds, with charged or polar R-groups, such as UroA. Compounds in this class are less lipophilic than ICZ and are more likely to form electrostatic interactions with conserved residues near the opening of the active site (in SRS regions 1-3; see Fig. S4), making it more difficult for them to access the heme center. Our analysis also predicts limited species-specific variability in the binding and recognition of UroA and ICZ by mouse and human forms of CYP1A1. The 3 amino acid differences found in the mouse CYP1A1 isoform (T111S, S116T, V228T (Fig. S4C-D), are inconsequential with respect to the positioning of functional, electrostatic or hydrophobic contacts in the CYP1A1 active site. While a large expansion in cavity volume was observed for the mCYP1A1 model (Fig. 5C-D), key hydrophobic and electrostatic contacts in SRS regions 1-6 were not disrupted.

### In vitro CYP1A1/1B1 metabolism of ICZ

ICZ modeling data provided us with the likely site of hydroxylation of ICZ at the C2 position. This allowed a synthesis scheme to be devised to synthesize the appropriate ICZ metabolites. Because ICZ is a symmetric molecule that may be hydroxylated at two sites, we synthesized both a mono- and di-hydroxylated ICZ derivative. LC-MS/MS and NMR analysis confirmed the identity of 5H, 11H-indolo[3,2b]carbazole-2,8-diol (2,8-dihydroxy-ICZ) and 5H, 11H-indolo[3,2b]carbazol-2-ol (2-hydroxy-ICZ) (Fig.S8 and S9). Using these compounds as standards we examined the ability of CYP1A1 to metabolize ICZ in a P450 microsomal assay system. Metabolism resulted in the formation of a peak at 7.9 min with the parent ion of 289.09 that match 2,8-dihydroxy-ICZ standard (Fig. 6A-C). Under the conditions of the assay 2-hydroxy-ICZ was not detected. Coincubation of UroA with ICZ in the P450 microsomal assay led to inhibition of ICZ metabolism to 2,8-dihydroxy-ICZ (Fig. 6D). In addition, ICZ can inhibit UroA metabolism to UroC (Fig. 6E-F).

**Figure 6.**
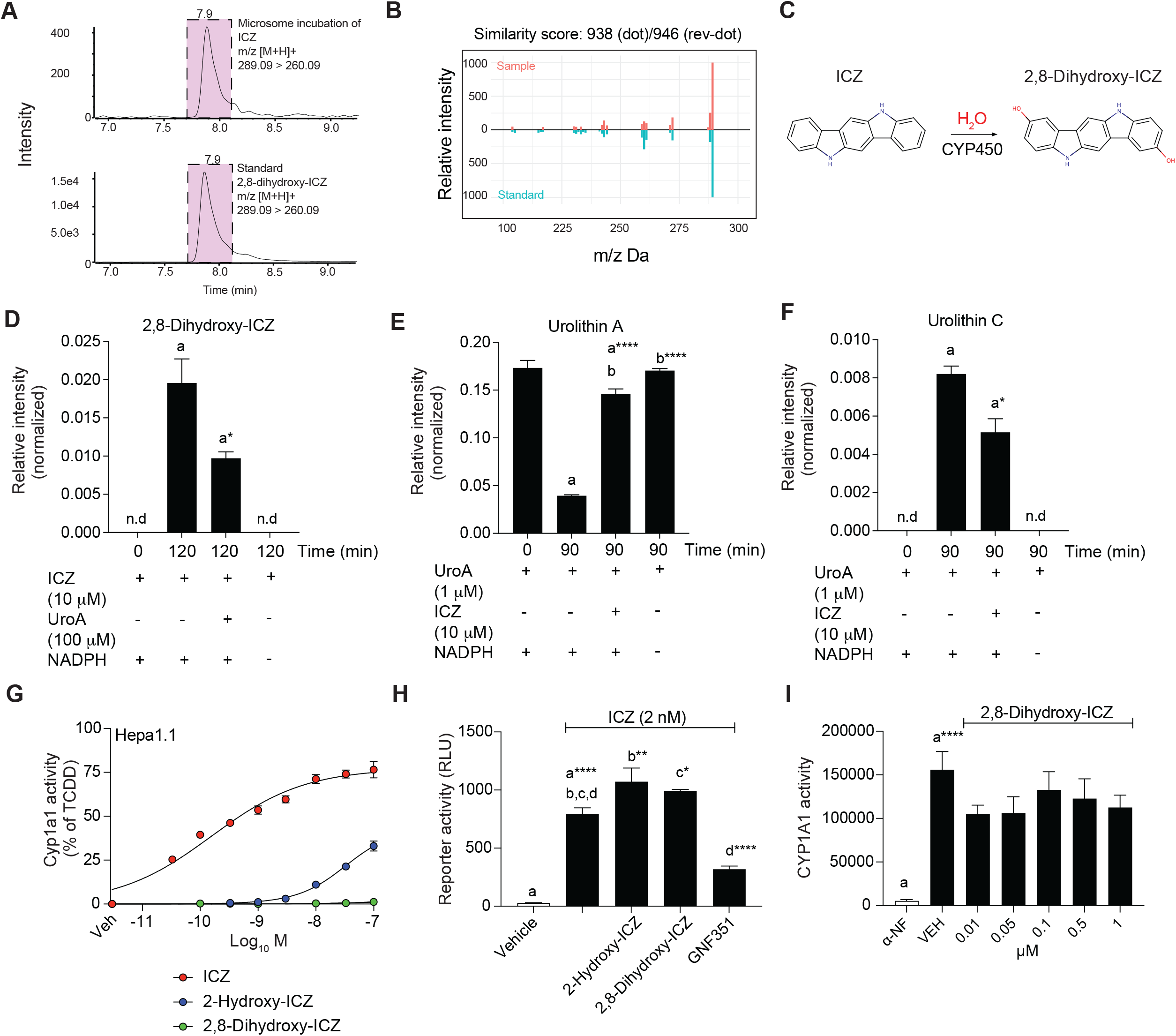
CYP1A1/1B1 metabolism of ICZ. (A) In vitro microsomal metabolism of ICZ results in the formation of 2,8-dihydroxy-ICZ. (B) MS/MS spectral similarity score confirms the identity of 2,8-dihydroxy-ICZ. (C) Scheme depicting the CYP1A1/1B1 metabolism of ICZ. (D) Urolithin A is capable of attenuating ICZ metabolism to 2,8-dihydroxy-ICZ in a microsomal metabolism assay. (E) ICZ can inhibit the metabolism of UroA in a microsomal metabolism assay. (F) ICZ can inhibit the formation of UroC from UroA in a microsomal metabolism assay. Statistical difference was analyzed using two-tailed unpaired Student’s t tests. Data are presented as mean + SEM (error bars). *p< 0.05, ****p<0.0001. (G) *Cyp1a1* mRNA levels were assessed in Hepa 1.1 cell line treated with various concentrations of ICZ, 2-hydroxy-ICZ, and 2,8-dihydroxy-ICZ. (H) Hepa 1.1 cells were treated with ICZ in the absence or presence of 1 µM 2-hydroxy-ICZ, 2,8-dihydroxy-ICZ, or the AHR antagonist GNF351 and AHR reporter activity assessed. (I) The in vitro microsomal assay was performed using Hepa 1.1 microsomes exposed to increasing concentrations of 2,8-dihydroxy-ICZ. Statistical difference was analyzed using one-way ANOVA with Tukey’s multiple comparisons. Data are presented as mean + SEM (error bars). *p< 0.05, **p< 0.01, ****p<0.0001.

### 2,8-Dihydroxy-ICZ fails to activate the AHR and is a poor CYP1A1 substrate/inhibitor

CYP1A1 metabolizes ICZ and likely attenuates ICZ-AHR signaling, thus setting up a negative feedback loop. To test this concept, we examined the ability of the downstream CYP1A1 metabolites of ICZ, 2-hydroxy-ICZ and 2,8-dihydroxy-ICZ, to activate the AHR in a cell-based AHR reporter cell line Hepa 1.1 assay. In comparison to ICZ, 2-hydroxy-ICZ exhibits ∼550-fold less activation potential based on ED_25_ dose (Fig. 6G), while 2,8-dihydroxy-ICZ minimal to no activation potential at the doses tested. Next, whether 2-hydroxy-ICZ and 2,8-dihydroxICZ can antagonize ICZ mediated AHR activation was tested, and neither ICZ metabolite was able to significantly antagonize ICZ (Fig. 6H). In contrast, the antagonist N-(2-(1H-indol-3-yl)ethyl)-9-isopropyl-2-(5-methylpyridin-3-yl)-9H-purin-6-amine (GNF351) effectively antagonized ICZ. We have shown that ICZ is readily metabolized by CYP1A1, thus whether 2,8- dihydroxy-ICZ is also a substrate or inhibitor of CYP1A1 was tested. In the P450 microsomal assay 2,8-dihydroxy-ICZ failed to exhibit a dose-dependent inhibition of CYP1A1 activity (Fig. 6I). Thus, 2,8-dihydroxy-ICZ appears to be the terminal CYP1A1 metabolite and is neither an AHR ligand nor CYP1A1 substrate. These data establish the AHR dependent expression and metabolic potential of CYP1A1 as a potent negative feedback regulator of ICZ-mediated AHR activity.

### Support for the ICZ-AHR-gut-lung axis

We wanted to further explore the hypothesis that ICZ enters the lymphatic system leading to enhanced AHR activation in the heart and lungs. *Cyp1a1* expression in the duodenum, liver and lung was assessed in mice on a chow diet. Comparison of lung and duodenal Cyp1a1 expression within individual mice revealed a significant positive correlation (Fig. 7A). In contrast, comparison of individual liver and duodenal Cyp1a1 expression within the treatment group did not significantly correlate (Fig. 7B). Next, we utilized the surfactant Pluronic L-81 previously demonstrated to disrupt chylomicron formation in the intestinal tract and thus uptake into the lymphatic system.^24, 25^ Mice were orally administered 20 mg of broccoli slurry with or without Pluronic L-81, after 12 h tissues isolated and *Cyp1a1* mRNA levels were assessed. The ratio of hepatic/pulmonary *Cyp1a1* revealed that Pluronic L-81 increased the hepatic/pulmonary Cyp*1a1* ratio (Fig. 7C). This data set was also plotted on an individual mouse basis and demonstrated that Pluronic L-81 decreased broccoli-mediated pulmonary *Cyp1a1* expression (Fig. 7D). This data further supports the concept that ICZ is entering the lymphatic system.

**Figure 7.**
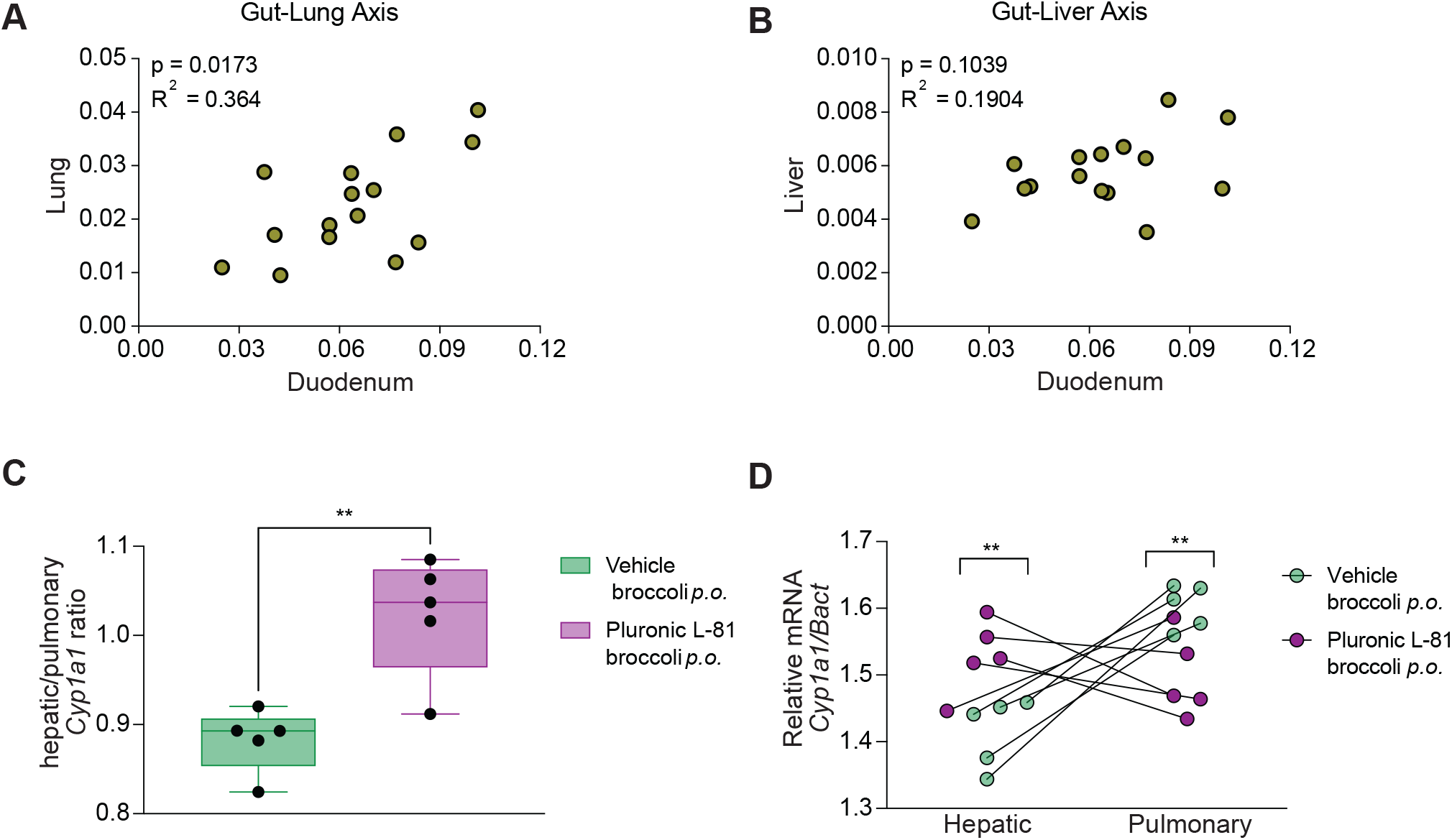
Assessment of possible lymphatic spread of an AHR ligand. (A) Significant correlation of pulmonary *Cyp1a1* duodenal *Cyp1a1* mRNA levels in C57BL6/J mice on a chow diet. (B) Lack of correlation between hepatic and duodenal *Cyp1a1* mRNA levels in C57BL6/J mice on a chow diet. (C) Pluronic L-81 increases the ratio of hepatic/pulmonary *Cyp1a1* upon co-treatment upon oral administration of a broccoli slurry. (D) The relationship between hepatic and pulmonary Cyp1a1 expression in individual mice from panel C. Statistical difference was analyzed using unpaired Student’s *t*-test, *n* = 5/group. Mean hepatic/pulmonary *Cyp1a1* ratio ± SEM were calculated, and significance was assessed. Data are presented as mean + SEM (error bars).**p< 0.01. Pearson correlation coefficient analysis was utilized in panels A and B, n= 15.

## DISCUSSION

Plants have evolved to synthesize secondary metabolites to deter consumption by herbivores and omnivores. In parallel, such animals have adaptively coevolved detoxification strategies utilizing a broad family of cytochrome P-450 (CYP) enzymes to counteract xenobiotic secondary metabolite toxicity following dietary consumption. The substrate specificity of CYP varies widely as well as the tissue-specific pattern of expression, which in effect leads to the ability to metabolize a wide spectrum of phytochemicals and other xenobiotics. CYP1A1 substrate specificity includes many hydrophobic polycyclic aromatic hydrocarbons, flavones, and quinolines. CYP1A1 is expressed in barrier tissues such as skin, lung and intestinal tract and appears to be highest in the liver and lung. Relatively high-affinity CYP1A1 substrates in general contain 3 or more ring structures, typified by 7-ethoxyresorufin, many polycyclic aromatic hydrocarbons (e.g. benzo(a)pyrene), and a variety of naturally occurring flavones ^26^. These structural attributes are strikingly similar to high affinity AHR ligands. Thus, CYP1A1 can serve as a negative feedback loop to limit AHR activation^21^. The endogenous high-affinity ligand FICZ has been demonstrated to be subject to this feedback loop^27, 28^. Depending upon relative concentrations, a compound that exhibits weak AHR binding capacity and manifests as a CYP1A1 substrate or inhibitor could therefore increase the half-life of a more potent AHR ligand and lead to *Cyp1a1* induction. Indeed, this has been observed in cell culture experiments utilizing quercetin and resveratrol as CYP1A1 substrates and FICZ as a high-affinity AHR ligand^29^. In this report the concept of an AHR-ICZ negative feedback loop was confirmed by the observation that CYP1A1 metabolizes ICZ to 2,8-dihydroxy-ICZ, leading to inactivation of AHR activation potential, and thus completion of the negative AHR feedback loop.

In the intestinal tract, especially the duodenum, enterocytes express significant amounts of CYP1A1 that metabolically restricts systemic circulation of a specific subset of AHR ligands that also manifest as CYP1A1 substrates.^30^ This was first demonstrated in vivo by the observation that the deletion of ARNT expression from the intestinal epithelium leads to extra-intestinal AHR activation, likely due to a lack of intestinal CYP1A1 expression^31^. Conversely, constitutive expression of CYP1A1 in the intestinal epithelial cells leads to a depleted AHR activation potential, resulting in significant changes in type 3 innate lymphoid cells within the intestinal tract^22^. We have previously demonstrated in mice that the lung exhibits the highest level of constitutive CYP1A1 expression.^21^ In addition, lung also exhibited the highest level of AHR activation potential in organic solvent tissue extracts from *Ahr^-/-^* mice, thus indicating significant levels of AHR ligands.^21^ However, these results did not address whether lung AHR activation is from a locally produced AHR ligand or from systemic circulation. The presence of a proximal to distal basal (and inducible) *Cyp1a1* gradient within the small intestine would suggest that small intestinal CYP1A1 metabolism of broccoli-derived ICZ would restrict its systemic distribution.^30^ Such digestive metabolism is consistent with the protective role of the intestine to limit systemic exposure to potentially toxic dietary secondary metabolites. Conversely, loss of AHR and thus CYP1A1 expression within the duodenum of *Villin^Cre^Ahr^fl/fl^* mice appears to increase systemic ICZ distribution, as evidenced by lung *Cyp1a1* induction relative to control diet. A comparison between *Ahr^fl/fl^* and *Villin^Cre^Ahr^fl/fl^* mice exposed to a broccoli diet reveals similar levels of hepatic *Cyp1a1* expression, suggesting equivalent systemic ICZ delivery to the liver. In contrast, a non-equivalent *Cyp1a1* induction is observed within the lung, with the absence of intestinal AHR expression resulting in enhanced pulmonary *Cyp1a1* expression relative to controls.

The presence of UroA in a broccoli diet yielded a similar *Cyp1a1* tissue-specific induction pattern as that observed in mice without intestinal CYP1A1 expression. Indeed, the addition of UroA to a broccoli diet selectively increases lung and heart AHR activation, while no effect on AHR activation in the liver was observed. A likely explanation for the tissue-specific AHR activation pattern observed upon consumption of UroA and ICZ is that UroA acts as a competitive CYP1A1 substrate in the intestinal tract and allows a proportion of ICZ to “escape” resulting in systemic circulation. This is supported by the enhanced *Cyp1a1* induction observed in the lung in mice that do not express AHR in the intestinal epithelium. However, this does not explain the lack of induction in the liver. Considering the hydrophobicity of ICZ, it is likely that some of this compound partitions into chylomicrons within enterocytes that are subsequently transported into the lymphatic system, thus bypassing the liver. The lymphatic system delivers its contents into the subclavian vein and enters the heart, followed by pulmonary circulation. This concept is consistent with other reports of a gut-lung axis.^32^ Interestingly, the second highest constitutive *Cyp1a1* mRNA level among the tissues examined was observed in the heart ^21^. Indeed, pharmaceutical companies have designed drugs that when taken orally directly enter the lymphatic system, bypassing liver metabolism prior to systemic circulation.^33^ However, additional studies will be needed to firmly establish the role of the lymphatic system in the observed ICZ/gut/lung phenotype. The ability of UroA to increase duodenal and lung activation when mice are on a semi-purified diet was unexpected. One possible explanation is that there are ligands that are generated but rapidly metabolized in the absence of phytochemicals. Clearly, additional studies are also needed to address the mechanism of this observation. Our data support the concept of unequal tissue biodistribution of ICZ in mice, which we demonstrate is likely a consequence of enhanced “escape” from diminished digestive CYP1A1 metabolism. This would facilitate increased lymphatic ICZ uptake from the intestine and transport via the thoracic duct directly to the lung, thus bypassing portal transport and hepatic first pass metabolism.

The ligand specificity and activation potential of the hAHR varies considerably from the mAHR. Xenobiotics such as TCDD and PAHs have about 10-fold higher affinity for the mAHR versus hAHR.^34^ In contrast, many tryptophan metabolites such as indirubin, indole, 3-methyl indole and kynurenic acid have significantly higher activation potential for the hAHR compared to the mAHR.^35^ Urolithins are hAHR antagonists, yet fail to bind to the mAHR.^16^ That would imply that in mice urolithins are CYP1A1 substrates and not AHR ligands. In contrast, in humans the influence of urolithins on AHR activity would likely be more complicated. Thus, in humans the presence of ICZ and urolithins would be more difficult to predict and may well be dependent on whether urolithins exhibit higher affinity as CYP1A1 competitive substrates compared to AHR antagonist potential. Indeed, considering the low nM AHR activation potential and affinity of ICZ in humans, it is likely that urolithins would function more as mediators of increased ICZ half-life.^36, 37^ The CYP1A1 modeling data presented here support the concept that ICZ is a better substrate for CYP1A1 than UroA. Nevertheless, the more abundant UroA could still effectively compete with the low level of ICZ produced from broccoli consumption. Finally, in a complex mixture of phytochemicals from a plant diet it will be difficult to predict the level of systemic or even intestinal AHR activation.

## Significance

Data presented herein demonstrate the existence of a diet-gut-lung AHR axis that facilitates the transport of a human relevant diet-derived plant metabolite (ICZ) directly to the lung, bypassing the highly efficient intestinal-portal-hepatic P450 metabolic system. Furthermore, intestinal/lymphatic escape and transport along this axis can be amplified through combinatorial exposure to multiple plant metabolites (as would be present in a complex diet) that present as competitive CYP1A1/1B1 substrates within the intestinal tract. Significantly, increased pulmonary bioavailability of ICZ promotes a lung-specific increase in AHR transcriptional activity. It is important to note that AHR activation within barrier tissues is associated with increased epithelial resistance to exogenous stress (e.g., physical/chemical trauma and microbial pathology). Thus, reduced intestinal metabolism and redirection of ICZ and other hydrophobic dietary AHR ligands to the lungs may provide a protective health benefit in this important barrier tissue. Finally, this report will lead to a reassessment of the dynamics of distribution of other hydrophobic chemicals present in the diet.

## METHODS

### Materials

UroA was synthesized as described in the supplemental information. All other urolithins were kindly provided by Francisco A. Tomás-Barberán (CEBAS-CSIC, Spain). α-naphthoflavone (α-NF) was purchased from Indofine Chemical Company (Hillsborough, NJ). TCDD was obtained from Dr. Stephen Safe (Texas A&M University). Pluronic L-81 was provided by Patrick Tso (University of Cincinnati). The broccoli cultivar Imperial was a provided by John Esslinger and Brian Campbell, and subsequently freeze dried, ground to a powder and stored at -80C. ICZ was obtained from Matrix Scientific (Columbus, SC) and GNF351 was synthesized as previously described.^38^ 5H, 11H-indolo[3,2-b]carbazol-2-ol (2-hydroxy-ICZ) and 5H, 11H-indolo[3,2-b]carbazole-2,8-diol (2,8-dihydroxy-ICZ) were synthesized as described here.

### Broccoli glucobrassicin quantification

The concentration of glucobrassicin in the broccoli cultivar Imperial was determined and was essentially the same as in the previously examined cultivar Lieutenant (1.6 µmoles/g dry weight).^14^

### Mouse studies

All mouse experiments were conducted at The Pennsylvania State University, University Park campus. All procedures were performed with approval and under the support of the Institutional Animal Care and Use Committee at The Pennsylvania State University. C57BL6/J mice were obtained from Jackson Laboratory (Bar Harbor, ME), congenic *Ahr^fl/fl^*, with the *Ahr*^b^ allele and *Villin^Cre^Ahr^fl/fl^* mice were bred in-house and maintained on a chow diet.

C57BL6/J mice were obtained from Jackson Laboratory (Bar Harbor, ME), bred in-house and maintained on a chow diet. Mice were placed on a semi-purified AIN-93G diet (Dyets Inc., Bethlehem, PA) for seven days prior to the initiation of experimental treatments. The broccoli cultivar Imperial was kindly supplied by John Esslinger and Brian Campbell, and was stored at -80°C, subsequently freeze-dried, and ground into a powder. AIN-93G diet was ground to a fine powder in a blender and mixed with 10% freeze-dried broccoli. UroA was added at 4 mg/g of diet where appropriate. Congenic *Ahr^fl/fl^* and *Villin^Cre^Ahr^fl/fl^* mice on a C57BL6/J background were generated as previously described.^30^

### Pluronic L-81 treatment

Female C57BL6/J mice were maintained with *ad libitum* access to a semi-synthetic diet (AIN93G) and water for 1 week. On day of treatment, mice were food restricted for 3 h (18:00-21:00 h) prior to oral administration of either 0.2 ml water (vehicle) or 5% (v/v) Pluronic L-81. After 30 min, mice were orally administered with 0.2 ml broccoli/water slurry (100 mg/ml freeze-dried broccoli) or 0.2 ml broccoli/water slurry supplemented with 5% Pluronic L-81. Mice were immediately returned to cages with over-night *ad libitum* access to semi-synthetic diet (AIN93G) and water. At 10:00 h, mice were euthanized and indicated tissues harvested for subsequent gene expression analysis.

### Cell culture

Hepa 1 and Caco2 cells were obtained from American Type Culture Collection and maintained in α-minimal essential medium (Sigma-Aldrich, St. Louis, MO) supplemented with 10% or 15% fetal bovine serum (Gemni Bio-Products, West Sacramento, CA), respectively, 100 U/ml penicillin and 100 µg/ml streptomycin. Cells were maintained at 37°C in a humidified incubator in 95% air and 5% CO_2_.

### CYP1A1/1B1 activity assay

Hepa1 cells were treated with 5 nM TCDD for 24 h to increase CYP1A1 protein levels. Microsomes were isolated from cells washed with phosphate-buffered saline, trypsinized and pelleted by centrifugation at 100 × g for 3 min. The cell pellet was then washed with phosphate-buffered saline, pelleted, and resuspended in 0.25 M sucrose, 10 mM Tris-HCl (pH 7.5), with protease inhibitors. The cells were manually homogenized in a Dura-Grind Dounce-Type homogenizer (Wheaton). The cell homogenate was centrifuged at 10,000 × g for 10 min at 4°C and the supernatant transferred and centrifuged at 42,000 × g for 90 min at 4°C. The resulting microsomal pellet was resuspended in the homogenization buffer and the protein concentration was determined. Microsomes were stored at -80°C for CYP1A1 activity assay. The P450-Glo CYP1A1/1B1 assay kit and NADPH regeneration system and luciferase assay kit were purchased from Promega (Promega, Madison, WI) and utilized under the following conditions: 12.5 μL of CYP reaction mixture including microsomes, Luciferin-CEE and potassium phosphate buffer, and 12.5 μL of test compound in distilled water were combined and pre-incubated at 37°C for 10 min in white 96-well plate. The reactions were initiated by adding an equal volume of NADPH regeneration system (25 μL) and placed at 37°C for 25 min. A final volume 50 µL of reactions were performed in triplicate and contained 10 μg of isolated microsomes in 100 mM K_2_PO_4_ (pH 7.4) buffer, 30 μM Luciferin-CEE, test compound, and NADPH regeneration system. Reactions were terminated by the addition of 50 μL of luciferin detection reagent and incubated at 37°C for 10 min. Luminescence was measured using a luminometer, and a control without microsomes was used to measure background. Relative CYP1A1 activity was expressed as the percentage of the luminescence after incubation with vehicle.

### CYP1A1/1B1 assays coupled to MS analyses

40 µg microsomes isolated from Hepa1 cells treated with 5 nM TCDD for 24 h, or lung microsomes (100 µg) from mice on the semi-purified diet, were incubated with various compounds and incubation times as indicated. The reactions were carried out at 37 °C in an incubator in the presence of NADPH regeneration system and terminated by addition of 800 µL ice cold 100% methanol (v/v) containing 1 µM chlorpropamide as internal standard. Then mixtures were kept in -20 °C for 30 mins followed by centrifugation at 12,000 × g for 20 min at 4 °C. The supernatants were dried under vacuum and reconstituted in 60 μL 50% methanol (v/v). After centrifugation, supernatants were transferred to autosampler vials for LC-MS/MS analysis as described in general chemistry methods section.

### RNA isolation and real-time quantitative PCR

Tissues were snap frozen in liquid nitrogen, placed in TRI Reagent (Sigma) and homogenized (6,500 rpm, 30 s) with 10-15 1 mm silica/zirconia beads (Biospec Products) using a BeadBlaster-24 homogenizer (Benchmark). Total RNA was initially isolated using TRI Reagent (Sigma-Aldrich) and then re-purified using RNeasy columns (Qiagen) following manufacturers’ protocols. Purified total RNA was used to generate cDNA using High-Capacity cDNA reverse transcriptase kit (Applied Biosystems) following manufacturers’ protocol. Quantitative real-time PCR was performed using primers (Table S1) as previously described^14^. Quantitative real-time PCR was performed using Perfecta SYBR green (QuantaBio) and the following cycling conditions (95°C, 3 min; 40 cycles: 95°C, 30 s; 58°C, 20 s, 72°C, 45 s; 72°C, 2 min). Melt-curve analysis revealed a single peak consistent with a single product.

### Capillary immunoblotting analysis

Microsomes were diluted to 0.5 mg/mL in 0.1X Sample Buffer (Protein Simple, San Jose, CA) and subjected to WES^TM^ capillary system analysis. The primary antibody for CYP1A1 (Proteintech, 13241, 1:100) was visualized with the WES anti-rabbit secondary HRP-conjugated antibody (Protein Simple). As a loading control for normalization, total protein was determined using Total Protein Detection Module (protein Simple, DM-TP01). For quantification, the chemiluminescence peak area of CYP1A1 was normalized to the total protein peak area for each sample.

### Computational Docking Analysis

The ligand binding affinity and target oxidation sites for UroA and ICZ were investigated using Autodock 4.2 and Autodock Vina, for both human and mouse models of CYP1A1^39–41^. The crystal structure of CYP1A1 [4I8V; copy A^42^] was used to prepare a docking model for the human enzyme, and as a template for a mouse homology model, which was developed using the SWISS-MODEL server^43^. All docking models were validated for overall quality, clashes and outliers using SAVESv6.0 server (https://saves.mbi.ucla.edu/)44-46 and ChimeraX^47^ using the ISOLDE plugin^48^. Crystal structure conformational bias was also reduced in substrate-free, heme-bound homology model of mouse CYP1A1 via MD simulation, using a simulated annealing and energy minimization routine (CHARMM36 force field) in VMD (QwikMD (NAMD); advanced run; standard settings)^49–51^.

Autodock 4.2 was run using standard settings, using the Genetic Algorithm search parameter (long evaluations, 20-100 GA runs) as previously described^52^. Grid box parameters were generated in Autogrid 4 for both Autodock 4.2 and Autodock Vina, using a 60-80 Å3 grid centered on the heme center. Docking results were analyzed using Autodock 4.2, The PyMOL Molecular Graphics System, Version 2.5.2 (Schrödinger, LLC) and LigRMSD 1.0^53, 54^. Active site cavity volumes of docking models were analyzed and compared using CAVER 3.0^55^.

### Quantification and Statistical Analysis

Values were expressed as mean ± SEM. Data were compared using either Student’s *t* test or one-way analysis of variance with Tukey multiple comparison post-test in GraphPad Prism 9.1.1 (GraphPad Software, Inc, La Jolla, CA, USA) to establish statistical significance between different groups. The value of *p* < 0.05 was considered statistically significant (\**p* < 0.05; \*\**p* < 0.01; \*\*\**p* < 0.001; \*\*\*\**p* < 0.0001). No data was excluded. All experiments were repeated at least twice.

## CHEMICAL SYNTHESIS

### Chemistry synthetic scheme

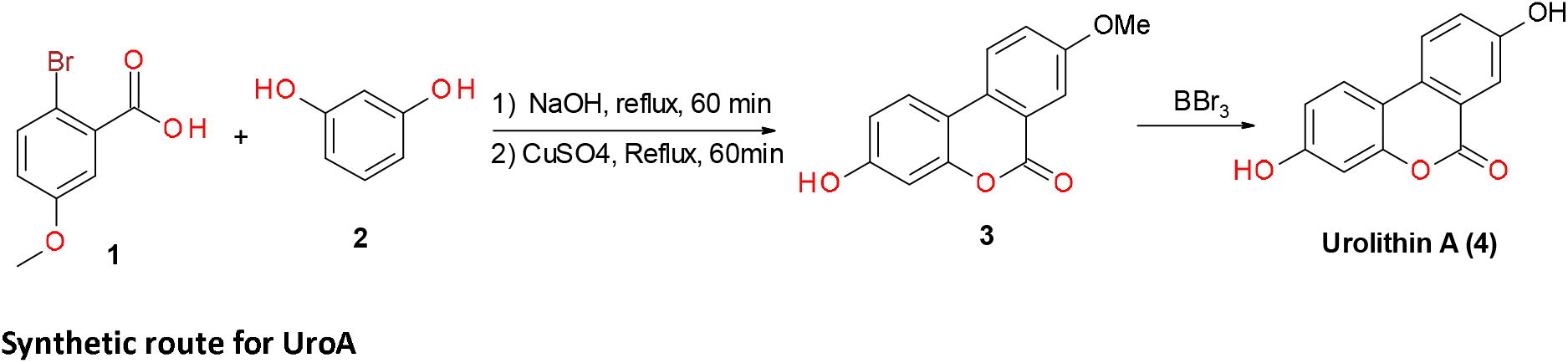

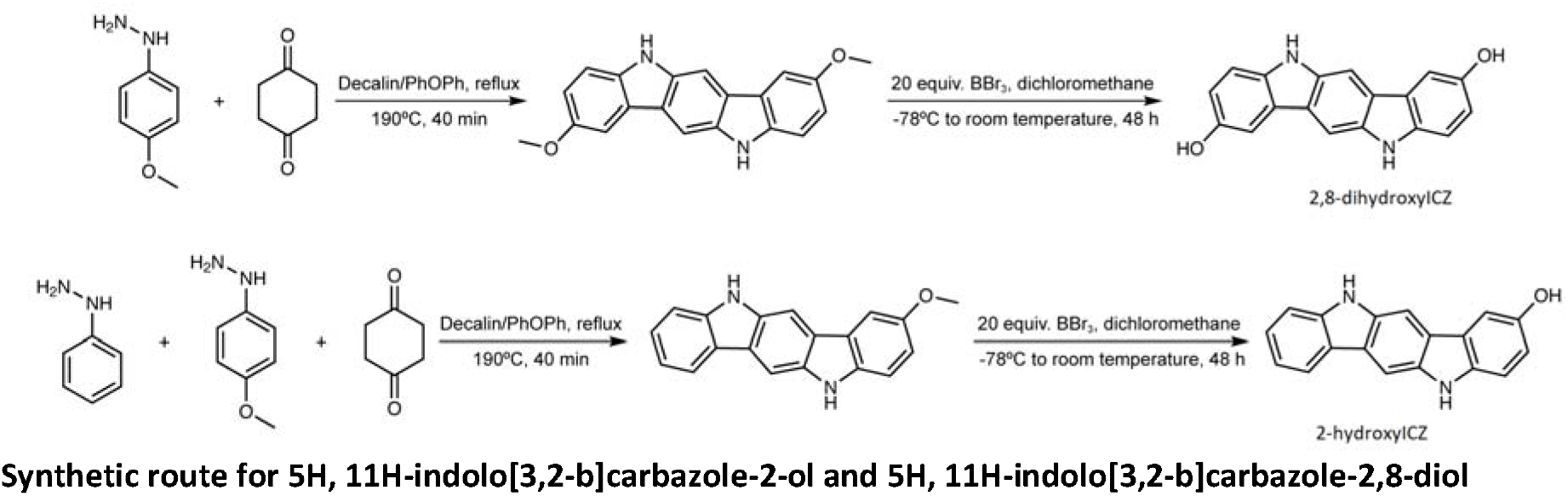

### General chemistry methods

LC/MS method section: Reagents and solvents were purchased from commercial vendors and were of the highest purity available and used without further purification unless otherwise noted. Qualitative/quantitative analysis was performed by reverse phase UHPLC using a Prominence 20 UFLCXR system (Shimadzu, Columbia, MD) with a Waters (Milford, MA) ACQUITY UPLC BEH C18 column (2.1 × 100 mm, 1.7 µm particle size) maintained at 55 °C and a 20 min aqueous acetonitrile gradient, at a flow rate of 250 µL/min. Solvent A was water with 0.1% formic acid and Solvent B was acetonitrile with 0.1% formic acid. The initial condition were 97% A and 3 % B, increasing to 45% B at 10 min, 75% B at 12 min where it was held at 75% B until 17.5 min before returning to the initial conditions. The eluate was delivered into an AB SCIEX TripleTOF™ 5600 System (QTOF) using a Duospray™ ion source (AB SCIEX, Framingham, MA). Purities of assayed compounds were in all cases greater than 95%, as determined by a Waters 2695 HPLC system (Waters, Milford, MA, U.S.A) with a Restek (Bellefonte, PA, U.S.A) HPLC C18 column (4.6 × 150 mm, 5 µm particle size) and a Viva C18 guard cartridge (10 × 4.0 mm, 5 µm particle size) with a 10 µL of injection volume. The mobile phase solvent A was 0.1% formic acid in water, and solvent B was 0.1% formic acid in acetonitrile. The gradient program was: 0-18 min, 5-45% B in A; 18-22 min, 45-90% B in A; 22-27 min, 90% B in A; 27-27.5 min, 90-5% B in A; 27.5-35 min, 5% B in A at a flow of 1 mL/min.

NMR method section: Each compound was dissolved in methanol-d4 or DMSO-d6. All NMR data were acquired at 298 K on Bruker Avance NEO 600 MHz spectrometer (Bruker Biospin, Rheinstetten, Germany) equipped with a 5 mm TCI cryoprobe. 1D ^1^H NMR spectra and a series of 2D NMR spectra, including ^1^H -^1^H TOCSY, ^1^H -^1^H COSY, ^1^H -^1^H JRes, ^1^H -^13^C HSQC, and ^1^H -^13^C HMBC, were recorded and processed as previously described with some modifications.^56^ The ^1^H and ^13^C chemical shifts were referenced to deuterated solvent residual proton signal at 3.31 ppm and ^13^C signal at 49.5 ppm in methanol-d4, respectively and at 2.50 ppm and 39.5 ppm in DMSO-d6, respectively. Coupling constants (J value) are reported in Hz. Spin multiplicities are described as s (singlet), br (broad singlet), d (doublet), t (triplet), and m (multiplet).

### Synthesis of UroA

**UroA (3,8-dihydroxy-6H-dibenzo[b,d]pyran-6-one, 4).** Urolithin A (**4**) was prepared in two steps starting from 2-bromo-5-methoxybenzoic acid (**1**) and resorcinol (**2**) as previously reported.^57^

#### Step 1

A mixture of 2-bromo-5-methoxybenzoic acid (**1**) (2.0 g; 9.3 mmol), Resorcinol (**2**) (2.05 g; 18.6 mmol), and sodium hydroxide (0.82 g; 20.46 mmol) in water (15 ml) was heated under reflux for 1 hour. After the addition of 10 % aqueous solution of CuSO_4_ (5 mL), the mixture was refluxed for additional 1 h. The mixture was allowed to cool to room temperature and the residue was filtered and washed with cold water to give 3-hydroxy-8-methoxy-6H-dibenzo [*b,d*] pyran-6-one (**3**) in 40% yield.

#### Step 2

To a suspension of **3** (0.60 g; 2.75 mmol) in dry CH_2_Cl_2_ (10 mL) at 0°C, was added 1 M solution of BBr_3_ in dry CH_2_Cl_2_ (8.25 mL, 8.25 mmol) over a period of 10 min. The mixture was left at 0°C for 1 h, allowed to warm up to room temperature and stirred for additional 17 hours. The reaction mixture was treated with ice and the resulted precipitate was filtered and washed with cold water to give a yellow solid which was heated to reflux in acetic acid for 3 h. The hot solution was filtered, and the residue was washed with acetic acid, then with diethyl ether to yield the title compound, UroA (**4)** as a yellow solid (85% yield). Structure of **4** was confirmed by ^1^H and ^13^C NMR. HRMS(m/z): [M-H]^-^ calcd for C_13_H_8_O_4_, 227.0344; found, 227.0347.

### General procedures for hydroxyindolo[3,2-*b*]carbazole synthesis

Synthetic routes were presented in scheme 1. All the reactions were performed in double-neck round bottom flasks. Purification of compounds were performed by flash column chromatography, which was packed by Synthware flash chromatography column (Kemtech America, inner diameter = 13.4 mm, length = 254 mm, outer diameter =17 mm) and silica gel (Sigma-Aldrich, pore size = 60 Å, 200-400 mesh particle size). The identity and purity of the compounds were determined by NMR and HPLC.

### Synthesis of 5H, 11H-indolo[3,2-b]carbazole-2,8-diol

#### Step 1

4-Methoxyphenylhydrazine hydrochloride (3.49 g, 20 mmol), 1,4-cyclohexanedione (1.12 g, 10 mmol), 12.5 mL decalin and 12.5 mL phenyl ether were added into a 100 mL round bottom flask. Subsequently, the mixture was stirred at 190°C for 40 min with reflux. The resulting mixture was cooled down to room temperature and then subjected to vacuum filtration to give a black solid. The solid was purified by flash column chromatography with isocratic elution of a mixture of hexane and ethyl acetate (v/v, 1/2) to afford 2,8-dimethoxyindolo[3,2-*b*]carbazole as a yellowish solid and changed to grey-greenish color gradually.

#### Step 2

20 equiv. boron tribromide (BBr_3_) was added dropwise into 20 mL dichloromethane (DCM) solution of 2,8-dimethoxyindolo[3,2-*b*]carbazole (100 mg) at -78°C (dry ice/acetone bath) under nitrogen. The dry ice/acetone bath was then removed, and the resulting mixture was stirred at 150 rpm and room temperature for 48 h. Ice/water was then added and the mixture extracted with ethanol (2 × 75 mL). The organic layer was dried with MgSO_4._ Then the solvent was removed by rotary evaporation, and the residue was purified by flash column chromatography with isocratic elution of a mixture of hexane and ethyl acetate (v/v, 1/2) to give 2,8-dihydroxyindolo[3,2-*b*]carbazole as a yellowish solid and slowly turned to grey-greenish color. Structure of synthesis was confirmed by NMR and HRMS respectively. ^1^H-NMR (600 MHz, DMSO-d6, 25°C): δ 10.49 (s, 2H, NH), 7.87 (s, 2H, =CH), 7.49 (d, *J* = 2.40 Hz, 2H, =CH), 7.25 (d, *J* = 8.50 Hz, 2H, =CH), 6.88 (dd, *J* = 8.50 Hz, 2.40 Hz, 2H, =CH); ^13^C-NMR (150 MHz, DMSO-d6, 25°C): δ 149.74, 135.59, 135.21, 123.28, 122.51, 114.45, 110.45, 104.74, 99.82. HRMS(m/z): [M+H]^+^ calculated for C_18_H_12_N_2_O_2_ was 289.0977, and determined to be 289.0940.

### Synthesis of 5H, 11H-indolo[3,2-b]carbazole-2-ol

By adding phenylhydrazine hydrochloride into the starting materials (4-methoxyphenylhydrazine hydrochloride (0.72 g, 5 mmol), phenylhydrazine hydrochloride (2.62 g, 15 mmol), 1,4-cyclohexanedione (1.12 g, 10 mmol), 12.5 mL decalin and 12.5 mL phenyl ether) and keeping the other methods the same as those in the synthesis of 2,8-dihydroxyindolo[3,2-*b*]carbazole, we successfully synthesized single hydroxyl substitute unsymmetrical ICZ derivative 2-hydroxyindolo[3,2-*b*]carbazole. The intermediate product 2-methoxyindolo[3,2-*b*]carbazole and the final product 2-dihydroxyindolo[3,2-*b*]carbazole were both grey-greenish solids eventually. Structure of synthesis was confirmed by NMR and HRMS respectively. ^1^H-NMR (600 MHz, DMSO-d6, 25°C): δ 10.9 (s, 1H, NH), 10.59 (s, 1H, NH), 8.82 (s, 1H, OH), 8.16 (m, 1H, =CH), 8.02 (s, 1H, =CH), 7.96 (s, 1H, =CH), 7.49 (d, *J* = 2.41 Hz, 1H, =CH), 7.43 (m, 1H, =CH), 7.35 (m, 1H, =CH), 7.25 (d, *J* = 8.49 Hz, 1H, =CH), 7.10 (m, 1H, =CH), 6.88 (dd, *J* = 8.49 Hz, 2.41 Hz, 1H, =CH); ^13^C-NMR (150 MHz, DMSO-d6, 25°C): δ 149.78, 141.02, 135.86, 135.16, 134.69, 125.06, 123.12, 122.58, 122.57, 122.27, 119.82, 117.25, 114.56, 110.55, 110.10, 104,81, 100.04, 99.95. HRMS(m/z): [M+H]^+^ calculated for C_18_H_12_N_2_O was 273.1027, and determined to be 273.0988.

## ACKNOWLEDGEMENTS

We thank John Esslinger and Brian Campbell for providing the broccoli used in this study. We also thank Juan Carlos Espin and Francisco Tomás for the series of urolithin compounds and Patrick Tso for generously providing the Pluronic L-81. We thank Marcia H. Perdew for critically reviewing this manuscript. This work was supported by the National Institutes of Health Grants ES028244 (GHP), ES028288 (ADP) and S10OD021750. This work was also supported by the USDA National Institute of Food and Federal Appropriations under Project PEN04607 and Accession number 1009993.

## AUTHOR CONTRIBUTIONS

G.H.P, C.M. designed the research. F.D., completed the in vitro microsomal assays and the qRT-PCR. D.C. performed the capillary immunoblotting analysis. I.A.M., completed the mouse experiments; A.A., performed the computational docking analysis; F.D., A.D.P., C.M., A.A., and I.A.M., contributed to data analysis. G.H.P., A.D.P., I.A.M., and A.A. wrote the manuscript. All authors discussed the results and commented on the manuscript.

## RESOURCE AVAILABILITY

### Lead contact

Further information and requests for resources and reagents should be directed to and will be fulfilled by the lead contact, Gary H. Perdew.

### Materials availability

Chemicals synthesized in this study will be made available for academic purposes upon request; commercial applications might require a contract and/or a completed Materials Transfer Agreement.

### Data and code availability

All date reported in this article will be shared by the lead contact upon request. The paper does not report original code. Any additional information required to reanalyze the data reported in this paper is available from the lead contact upon request. The metabolomics data is available at the NIH Common Fund’s National Metabolomics Data Repository (NMDR) website, the Metabolomics Workbench, https://www.metabolomicsworkbench.org where it has been assigned Study ID ST002481. The data can be accessed directly via its Project DOI: http://dx.doi.org/10.21228/M8H131.

