## Supplemental figures for "Complex chemical signals dictate Ah receptor activation through the gut-lung axis"

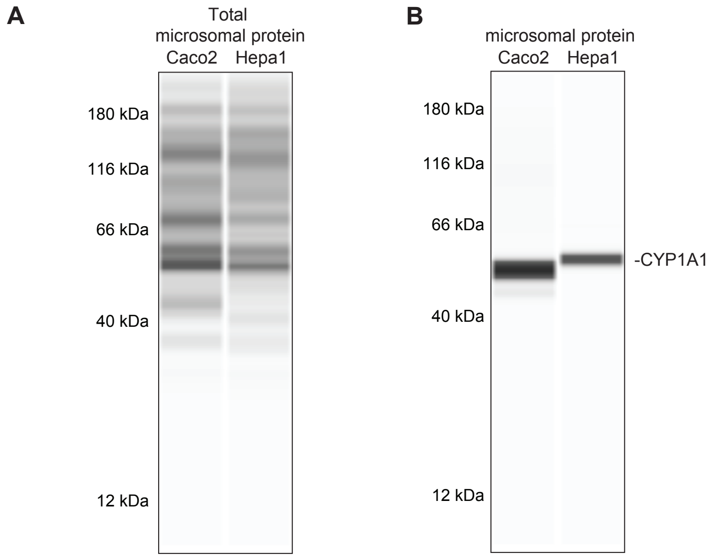


**Fig. S1.** TCDD induces CYP1A1 expression in Caco2 and Hepa 1 cells. The WES protein electrophoresis system was used to assess the presence of CYP1A1. (A) Total protein levels were examined in microsomes isolated from Caco2 and Hepa 1 cells treated with TCDD for 24 h. (B) An anti-CYP1A1 antibody was utilized to visualize the presence of CYP1A1 in microsomes.

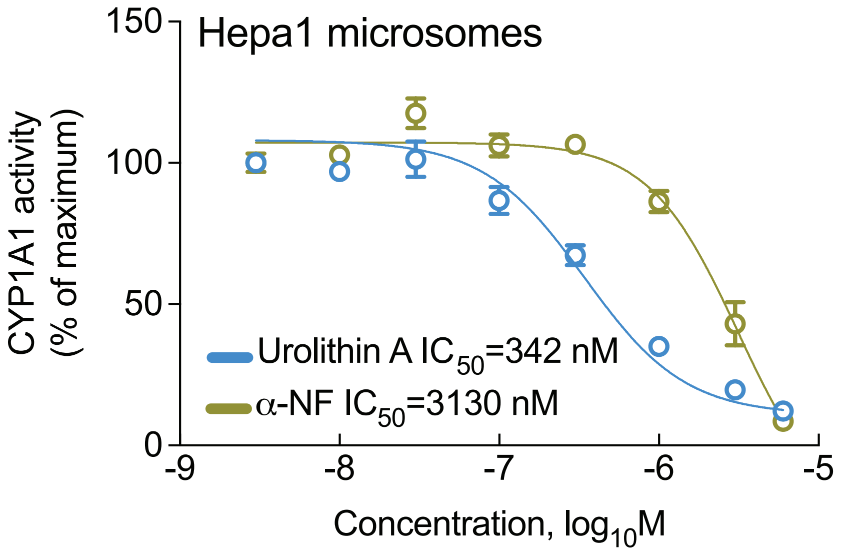


**Fig. S2**. Urolithin A and α-NF inhibit luciferin metabolism. Comparative analysis of dose-dependent inhibition of luciferin-CEE metabolism by UroA and α-NF in the Hepa 1 or Caco2 microsomal assay system.

.

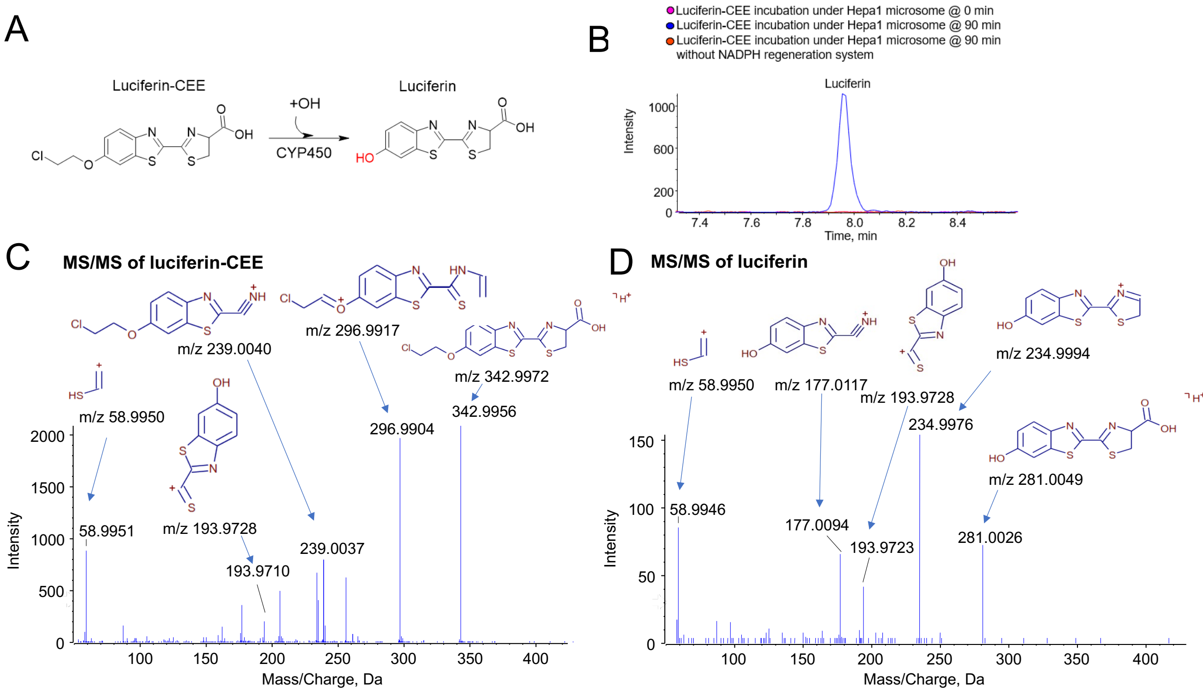


**Fig. S3.** Luciferin was produced from metabolism of luciferin-CEE incubated with microsomes isolated from Hepa1 cell.(A) Reaction route of luciferin-CEE being metabolized to luciferin. (B) Representative chromatographic spectrum of luciferin was observed by CYP1A1/1B1 incubation for 90 min, yet not present without NADPH regeneration system. (C) Fragment annotations of MS/MS of luciferin-CEE which was added in assay. (D) Fragment annotations of MS/MS of luciferin metabolized from luciferin-CEE in assay.

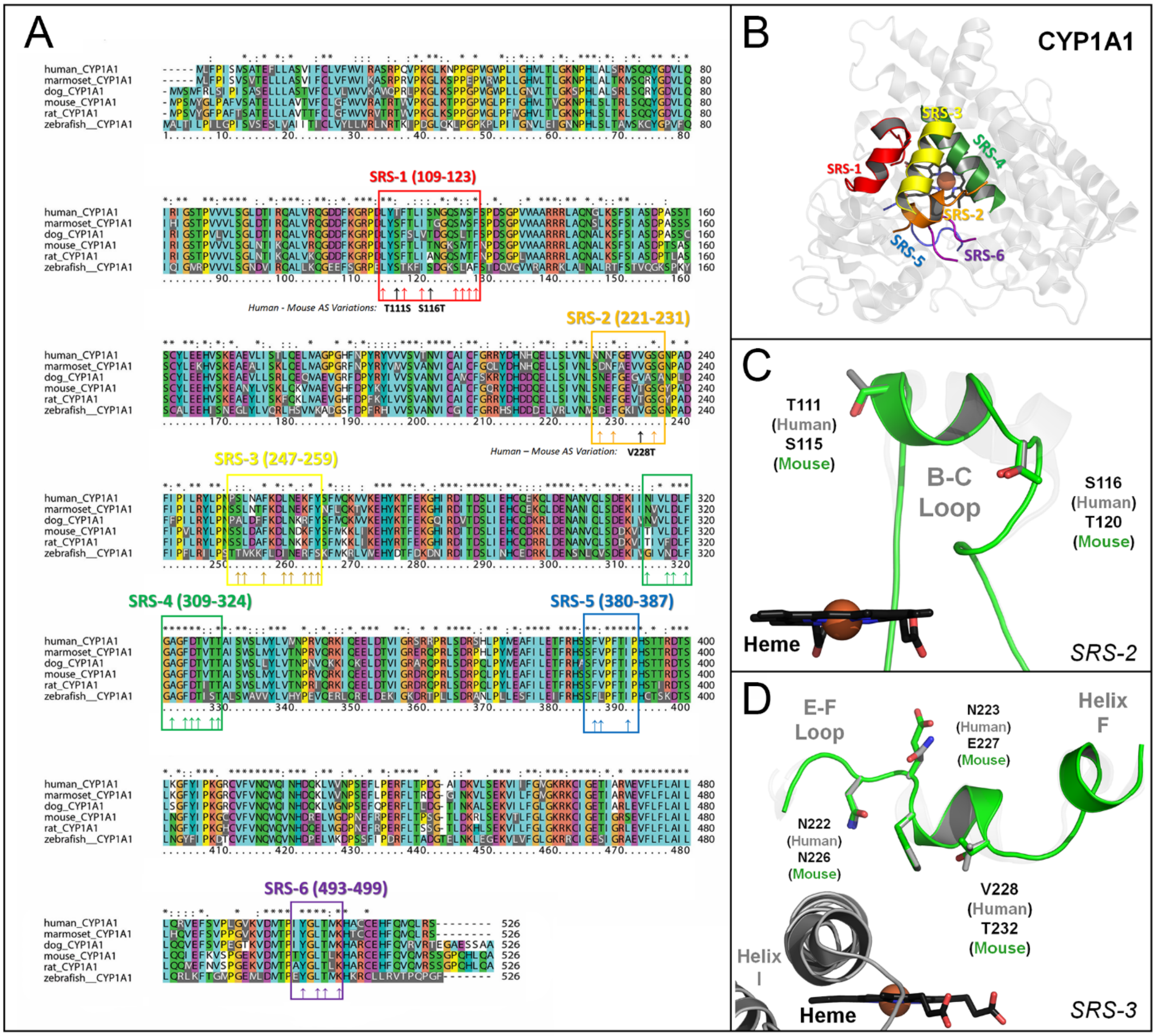


**Fig. S4.** CYP1A1 Substrate-Recognition Sequences and Structural Analysis. (A) Primary amino acid sequence alignment for reference CYP1A1 proteins from human, mouse, dog, rat, marmoset, and zebrafish are shown. Six Substrate-Recognition Sequences (SRS) motifs for CYP1A1 are highlighted: SRS-1 (109-123; red), SRS-2 (221-231; orange), SRS-3 (247-259; yellow), SRS-4 (309-324; green), SRS-5 (380-387; blue) and SRS-6 (493-499; purple). Amino acid numbering is for the human gene and well conserved residues oriented into the active-site pocket (in the crystal structure 4I8V) are highlighted with up arrows in each SRS region. Inward facing amino acids, that represent human to mouse active-site (AS) differences between human and mouse CYP1A1 (T111S, S116T, V228T) are further highlighted in bold, black text. (B) The position and interactions among the six SRS regions (SRS1-6) described above, are shown in 3D space, above the heme centered CYP active site, using the crystal structure of human CYP1A1 (4I8V; cartoon representation). (C) The mouse and human CYP1A1 active site are distinguished by only 3 primary amino acid differences. Two of these differences are found on the B'-helix, a part of the B-C helical loop in the SRS-2 motif. Mouse and human CYP1A1 exchange polar residues on either side of the B’-helix (T111 in human vs. S116 in mouse and S116 in humans vs. T120 in mouse). These sequence variations may change non-polar, hydrophobic contacts in the region, but are not predicted to alter electrostatic interactions significantly, as the polar, hydroxyl group of each residue remain essentially fixed in 3D modeling space. (D) The third key active site difference between mouse and human CYP1A1 is found in the SRS-2 region, which is comprised of the E-F helical loop and the F helix. Valine-228 (human) is replaced by a more polar, threonine molecule (T232 in mouse) which expands the active site cavity size (see Supplemental Figure 3). However, the overall topology of the SRS-2 motif remains highly similar in both mouse and human, as the conserved E-F loop asparagine (N222 (human) vs. N226 (mouse)) dominates substrate interactions in both human and mouse models. An additional, species-specific sequence variation in this region (N223 (humans) vs. E227 (mouse)) is also highlighted. While these residues are oriented out of the active site in the crystal structure of CYP1A1, it is possible that these residues can also orient into the active site in alternate, closed conformations of the CYP, also regulating species-specific differences in substrate metabolism.

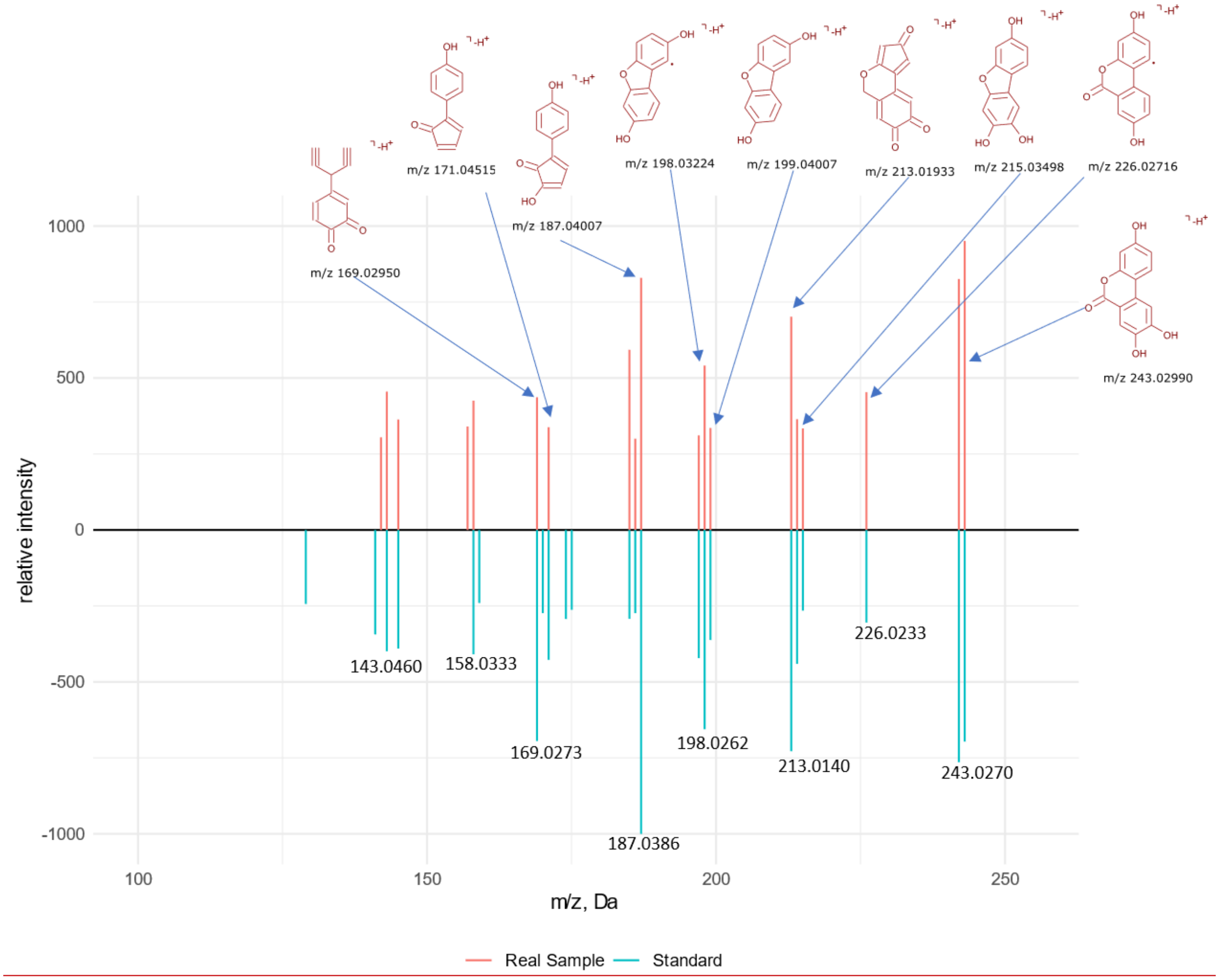


**Fig. S5.** Similarity plot of MS/MS data with fragment structures illustrates that within an in vitro microsomal incubation UroA is metabolized to UroC. LC-MS/MS validation of identity between authentic UroC and UroC detected in UroA incubation with microsomes isolated from Hepa1 cells.

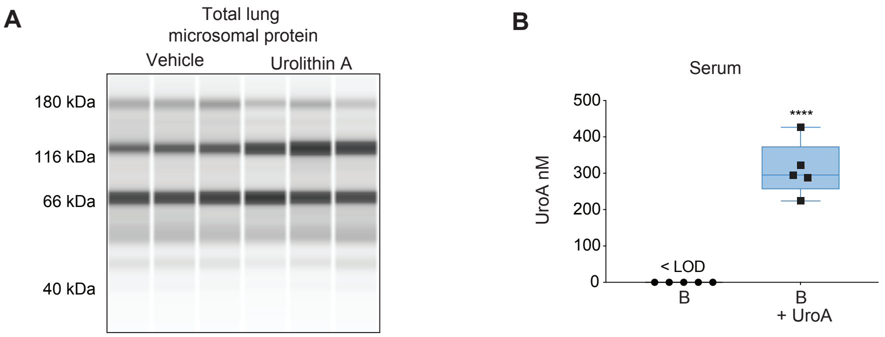


**Fig. S6.** Analysis of total protein in lung microsomes and Serum concentrations of UroA.
(A) Total protein determined by WES system from microsomes, control for CYP1A1 analysis. (B) UroA concentration in mouse serum collected from mice feed either a semi-purified diet or a semi-purified diet containing 4 mg of UroA/g of diet for 3 days.

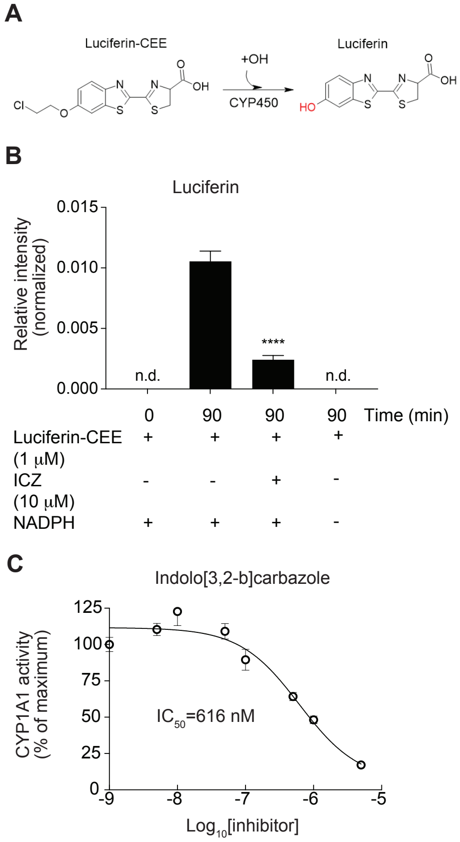


**Fig. S7.** ICZ inhibits CYP1A1 metabolism in a CYP1A1/1B1 microsomal assay. (A) Structural depiction of CYP1A1/1B1 metabolism of luciferin-CEE to luciferin. (B) ICZ significantly inhibits luciferin-CEE metabolism to luciferin in a Hepa 1 in vitro microsomal assays. (C) ICZ inhibits CYP1A1 metabolism of luciferin-CEE in a dose-dependent manner.

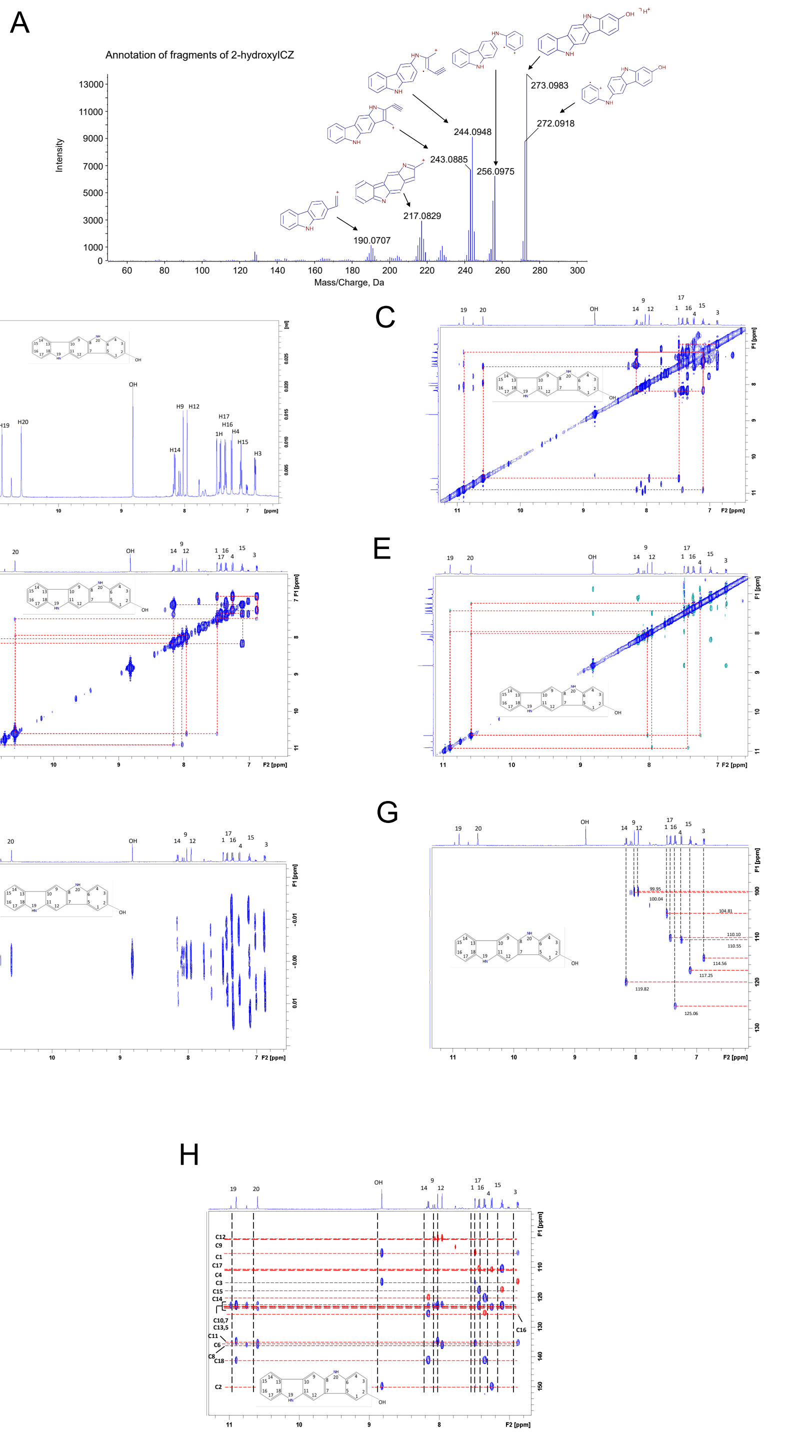


**Fig. S8.** Conformation of the structure of 2-hydroxyICZ by a combination of NMR and MS/MS. (A) Annotation of the MS/MS spectrum of 2-hydroxyICZ. (B) 1D $^{1}H$ spectrum of 2-hydroxyICZ with assignments. (C) $^{1}H$-$^{1}H$ TOCSY spectrum of 2-hydroxyICZ with signal assignments (in DMSO-$d_6$); (D) $^{1}H$-$^{1}H$ COSY spectrum of 2-hydroxyICZ with signal assignments (in DMSO). (E) $^{1}H$-$^{1}H$ ROESY spectrum of 2-hydroxyICZ with signal assignments (in DMSO). (F) $^{1}H$-$^{1}H$ JREs spectrum of 2-hydroxyICZ with signal assignments (in DMSO). (G) $^{1}H$-$^{13}C$ HSQC spectrum of 2-hydroxyICZ with signal assignments (in DMSO). (H) $^{1}H$-$^{13}C$ HMBC (blue) and HSQC (red) spectrum of 2-hydroxyICZ with signal assignments (in DMSO).

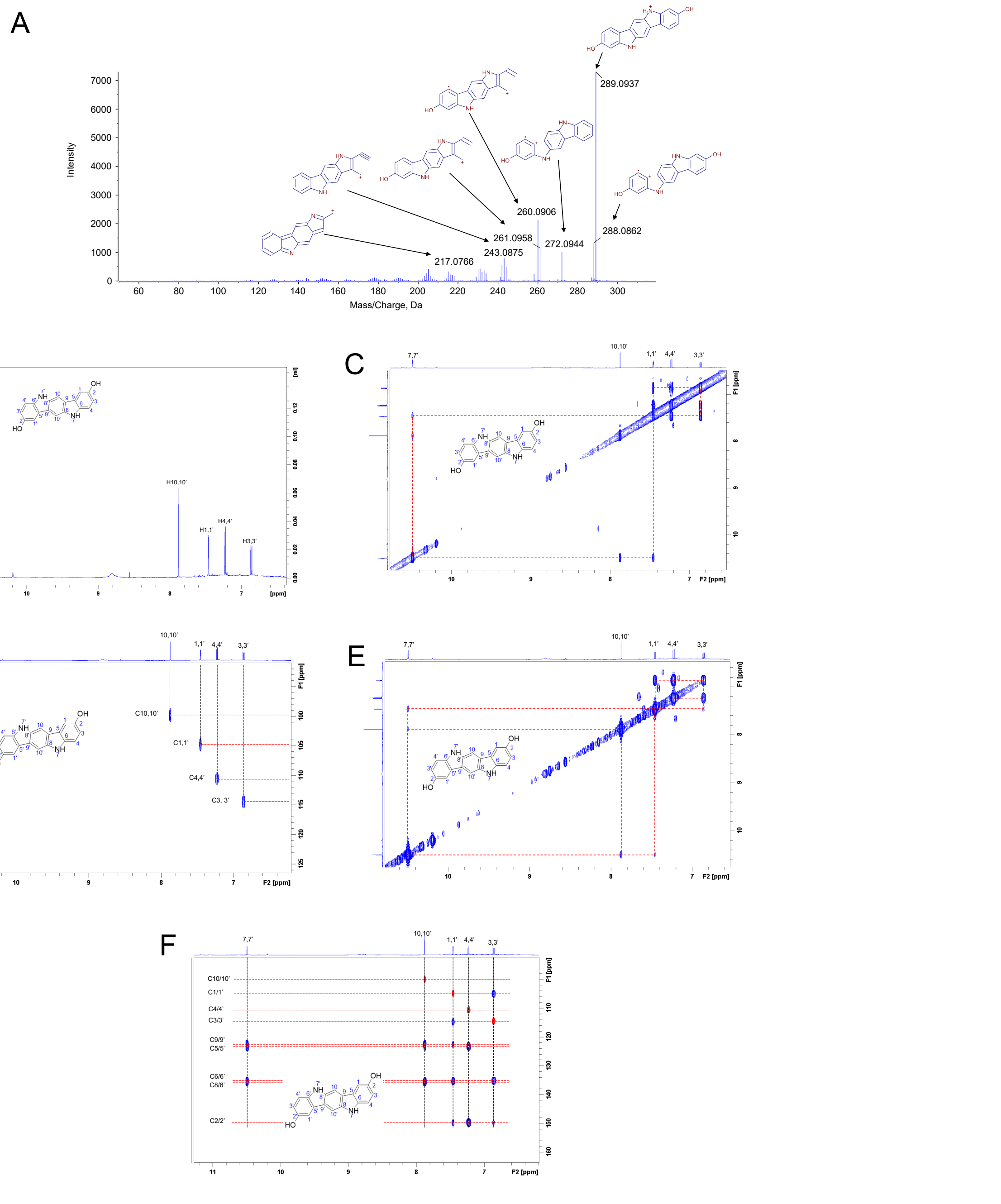


**Fig. S9.** Conformation of the structure of 2,8-hydroxyICZ by a combination of NMR and MS/MS. (A) Annotation of the MS/MS spectrum of 2,8-dihydroxyICZ. (B) 1D $^{1}H$ spectrum of 2,8-hydroxyICZ with assignments. (C) $^{1}H$-$^{1}H$ TOCSY spectrum of 2,8-hydroxyICZ with signal assignments. (D) 1H-$^{13}C$ HSQC spectrum of 2,8-dihydroxyICZ with assignments. (E) $^{1}H$-$^{1}H$ COSY spectrum of 2,8-dihydroxyICZ with signal assignments. (F) 1H-$^{13}$C HMBC (blue) and HSQC (red) spectrum of 2,8-hydroxyICZ with signal assignments.

**Supplementary Table S1. Computational Docking Analysis of Urolithin A and ICZ to human and mouse CYP1A1.**

| Substrate | | Computational Docking Program | | | | | |
|---|---|---|---|---|---|---|---|
| | | *Autodock 4.2* | | | | *Autodock Vina* | |
| | | **Human Cyp1A1** | | **Mouse CYP1A1** | | **Human Cyp1A1** | **Mouse CYP1A1** |
| | | Dissociation Constant[a] ($K_D$) in nM | Binding Energy[b] (kcal/mol) | Dissociation Constant ($K_D$) in nM | Binding Energy (kcal/mol) | Binding Energy[c] (kcal/mol) [$K_D$] | Binding Energy (kcal/mol) [$K_D$] |
| **Urolithin A** | *Max*[d] | **152** | **-9.3** | **191** | **-9.17** | **-10.9** *[17 nM]* | **-10.6** *[10 nM]* |
| | *Avg*[e] | *234 ± 85* | *-9.1 ± 0.2* | *308 ± 116* | *-8.9 ± 0.2* | *-10.5 ± 0.5* | *-10.2 ± 0.5* |
| **ICZ** | *Max* | **8** | **-11.1** | **84** | **-11.3** | **-14.1** *[49 pM]* | **-13.5** *[135 pM]* |
| | *Avg* | *8.0 ± 0.1* | *-11.1 ± .01* | *85.5 ± 2.1* | - | *-14.0 ± 0.2* | - |

[a]Dissociation binding constant (KD) were derived computationally using Autodock 4.2 analysis.

[b]Substrate Binding Energies were derived computationally using Autodock 4.2 analysis.

[c] Substrate Binding Energies were derived computationally using Autodock Vina. Because $K_D$ values are not computed manually in Autodock Vina, they were extrapolated to the Autodock 4.2 conversion scale using the following equation: $y = 0.5982\ln(x) - 12.304$, which was derived from Autodock 4.2 docking data presented in this table ($R^2 = 0.9994$).

[d]Max refers to the maximum affinity, or lowest-energy docking solution identified by either Autodock 4.2 or Autodock Vina. For Autodock 4.2 the solution selected from the top 20 docking conformations (long GA runs) and from an unlimited number of solutions in Autodock Vina.

[e]Avg refers to the average or cumulative dissociation constant $K_D$ and binding energy for all docking pose obtained for an individual substrate:model docking combination ($N < 20$). If an Avg value is not provided then the Max solution represented the only docking solution returned in the CYP active site.
